# Mechanosensation of the heart and gut elicits hypometabolism and vigilance in mice

**DOI:** 10.1101/2023.06.29.547073

**Authors:** Karen A. Scott, Yalun Tan, Dominique N. Johnson, Khalid Elsaafien, Caitlin Baumer-Harrison, Sophia A. Eikenberry, Jessica M. Sa, Guillaume de Lartigue, Annette D. de Kloet, Eric G. Krause

## Abstract

Interoception broadly refers to awareness of one’s internal milieu. Vagal sensory afferents monitor the internal milieu and maintain homeostasis by engaging brain circuits that alter physiology and behavior. While the importance of the body-to-brain communication that underlies interoception is implicit, the vagal afferents and corresponding brain circuits that shape perception of the viscera are largely unknown. Here, we use mice to parse neural circuits subserving interoception of the heart and gut. We determine vagal sensory afferents expressing the oxytocin receptor, hereafter referred to as NDG^Oxtr^, send projections to the aortic arch or stomach and duodenum with molecular and structural features indicative of mechanosensation. Chemogenetic excitation of NDG^Oxtr^ significantly decreases food and water consumption, and remarkably, produces a torpor-like phenotype characterized by reductions in cardiac output, body temperature, and energy expenditure. Chemogenetic excitation of NDG^Oxtr^ also creates patterns of brain activity associated with augmented hypothalamic-pituitary-adrenal axis activity and behavioral indices of vigilance. Recurrent excitation of NDG^Oxtr^ suppresses food intake and lowers body mass, indicating that mechanosensation of the heart and gut can exert enduring effects on energy balance. These findings suggest that the sensation of vascular stretch and gastrointestinal distention may have profound effects on whole body metabolism and mental health.

## Main

Perturbations to the internal milieu are relayed from peripheral organs to the brain to elicit compensatory responses that maintain homeostasis. For example, chemical and mechanical signals arising from the vasculature and gastrointestinal tract are transduced into action potentials carried by the vagus nerve into the brain via synapses in the nucleus of the solitary tract (NTS). The NTS integrates these interoceptive signals and routes them to effector brain nuclei that shape the sensation of thirst or hunger, and evoke compensatory responses aimed at mitigating cardiometabolic deficits^1^. Maintaining cardiometabolic homeostasis is fundamental to an organism’s vitality and requires integration of complex responses with autonomic, endocrine, and behavioral nodes. Given the diversity of these responses, it is perhaps not surprising, that the status of the internal milieu can profoundly influence the perception of the external environment. In this way, interoceptive signals originating from cardiovascular and gastrointestinal tissues may affect emotionality^2,3^ and cognition^4^ as well as cardiac output^5^ and nutrient selection^6^. Yet, relative to the sensory modalities that perceive the external environment, little is known about the processes by which an organism monitors sensations that originate from within itself.

Perturbations to the internal milieu impose stress on an organism, which we define as real or perceived threats to homeostasis. At a fundamental level, stress manifests from an organism’s perception that the energetic demands of its environment exceed metabolic fuel availability, and this spurs centrally-mediated responses that promote the mobilization and conservation of energy^7^. While interoception potently influences stress responding and the regulation of energy balance, the sensory modalities and brain circuits governing these processes are not well understood^8^. The internal milieu is sensed by vagal afferents, the soma of which, reside within the nodose ganglia. Studies by Kupari *et al* utilized single-cell RNA sequencing of nodose ganglionic neurons to identify gene clusters associated with specific groups of vagal neurons that may function as mechano-, chemo-, and/or nociceptors^9^. Building off this work, Bai *et al.*^10^ found that nodose ganglionic neurons that express mRNAs encoding for the oxytocin receptor (Oxtr) specifically function as mechanoreceptors that monitor stretch and tension exerted on the viscera. In the brain, Oxtr(s) are synthesized by neurons that mediate stress responding by altering the perception of emotionally salient stimuli in the external environment^11^. Conversely, the presence of Oxtr(s) within the nodose ganglia presents the possibility that these receptors also demarcate neurons that convey body-to-brain communication that initiates physiological and behavioral responses to stressors originating from the internal milieu.

Here, we use mice to test the overall hypothesis that NDG^Oxtr^ function as mechanosensors that sense force exerted on the viscera and recruit activation of brain nuclei orchestrating energy balance and stress responses. Towards this end, we reveal that NDG^Oxtr^ innervate the aortic arch and portions of the gastrointestinal tract, where they form structural endings consistent with mechanosensation. Fascinatingly, chemogenetic activation of NDG^Oxtr^ produced a phenotype resembling torpor, which is a state of hypometabolism that heterotherms enter when exposed to energetically demanding environments^12^. Collectively, our results demonstrate that perceived mechanosensation of the heart and gut potently alters metabolism and behavior to cope with threats originating from the internal milieu. The implication is that NDG^Oxtr^ can be studied to understand the etiology or reversal of a variety of stress-related diseases.

## RESULTS

### NDG^Oxtr^ comprise a subset of vagal neurons with structural endings that monitor stretch and tension exerted on the stomach, duodenum, and aortic arch

We visualized NDG^Oxtr^ by way of bilateral injection of Cre-inducible adenoassociated virus (AAV) that expresses the red fluorescent protein, mCherry, into the nodose ganglia of mice with Cre-recombinase directed to the *Oxtr* gene (i.e., Oxtr^Cre^ mice) (**Fig. 1a**). Three weeks later, mice were perfused and nodose ganglia were extracted and processed for RNAscope *in situ* hybridization for *Oxtr* mRNA followed by immunohistochemistry for NeuN (neuronal marker). Confocal imaging and analysis found that 83 ± 3.4% of mCherry labeled neurons contained *Oxtr* mRNA (**Fig. 1b-c**) indicating that Cre-recombination was directed to the *Oxtr* gene with the same high fidelity that we previously observed in the brain^13^. Conversely, 21 ± 1.8% of NeuN-labeled cells in the nodose ganglia expressed mCherry (**Fig. 1d-e**). These results are consistent with a prior study that found ≈20% of disassociated nodose ganglionic neurons were responsive to bath applied oxytocin^14^ and establish that Oxtr(s) are not ubiquitously expressed, but rather, delineate a subpopulation of nodose ganglionic neurons.

**Fig. 1.**
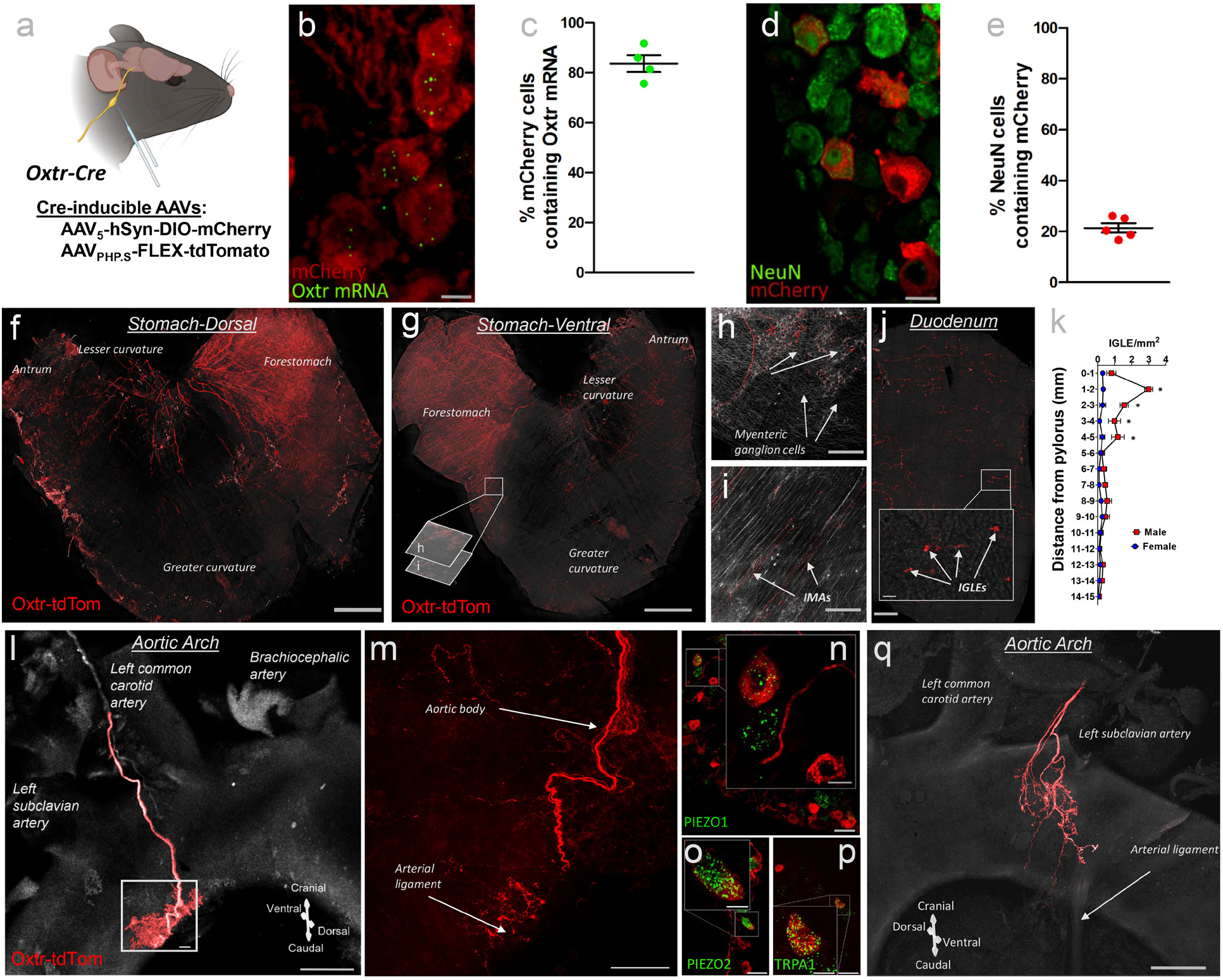
Subpopulations of neurons within the NDG ganglia contain Oxtr(s) and form structural endings in the stomach, duodenum and aortic arch that are likely to monitor tension and stretch. (**a**) Schematic illustrating application of AAV_5_-hSyn-DIO-mCherry (**b-e**) or AAV_PHP.S_-FLEX-tdTomato (**f-q**) bilaterally into the NDG. (**b-e**) Images and quantification of co-localization of mCherry and (**b, c**) *Oxtr* mRNAs or **(d, e**) NeuN in NDG^Oxtr^. (**f-k**) tdTomato fibers arising from NDG^Oxtr^ are found in the (**f-i**) stomach and (**j**) duodenum and form structural endings [IGLEs (**h, j_inset_**) and IMAs (**i**)] that are suggestive of their involvement in mechanosensation. Note that distance between sections in panels **h** and **i** is 100 µm. (**k**) Comparison of IGLE density across the length of the duodenum in males and females (n= 3 males, 3 females). (**l-m**) tdTomato fibers arising from NDG^Oxtr^ are also found in the aortic arch. (**n-p**) Co-localization of *Piezo1*, *Piezo2* and *Trpa1* mRNAs in tdTomato-labeled NDG^Oxtr^. (**q**) Similar innervation patterns in the aortic arch were observed in Oxtr^2A-cre^ knock-in mice (used in Bai *et al.)* delivered AAV_PHP.S_-FLEX-tdTomato, in NDG. Scale = 10 µm (**b, d, j_inset_, n_inset_-p_inset_**); 50 µm (**n-p**); 100 µm (**h-j, m**); 500 µm (**l, q)**; 1 mm (**f, g**). Bars represent SEM. *, P <0.05. Schematics made with Biorender.com.

To identify tissues innervated by NDG^Oxtr^, we administered Cre-inducible AAV that expresses the red fluorescent proteins, tdTomato, into the nodose ganglia of Oxtr^Cre^ mice. Two to three weeks later, mice were perfused and the nodose ganglia, gastrointestinal tract, and heart, including the aortic arch and aortic body were extracted. Organs were scanned as whole mount samples with a confocal microscope. As shown in **Fig. 1f-j**, tdTomato clearly labeled fibers within the stomach and duodenum and we sought to reveal whether such fibers exhibited terminal morphologies indicative of mechanoreceptors and/or chemoreceptors as previously described^15,16^. **Figures 1h-j** depict intramuscular arrays (IMAs) and intraganglionic laminar endings (IGLEs) with the latter juxtaposed to myenteric ganglion cells; however, mucosal endings innervating the intestinal villi were not observed. Prior studies determined that IGLEs function as mechanosensors detecting GI stretch^17-19^, thus, our results suggest that NDG^Oxtr^ monitor distention of the stomach and duodenum. Despite having similar numbers of tdTomato-labeled soma within the NDG, female mice exhibited significantly fewer tdTomato-labeled IGLEs, relative to males, and this effect was most pronounced in the proximal duodenum (**Fig. 1k**). Collectively, these results confirm and extend findings of another report indicating that NDG^Oxtr^ innervate the gastrointestinal tract where they function as mechanosensors that monitor stretch and tension exerted on the wall of the stomach and intestine^10^.

We also observed tdTomato-labeled fibers encapsulating a portion of the aortic arch **[Fig. 1l**]. These fibers formed a claw-like structure positioned between the left subclavian and left common arteries [Fig. 1m]. A prior study determined that this aortic-claw arises from nodose ganglionic neurons that express mechanically gated ion channels, PIEZO1/2, and function as mechanoreceptors, specifically arterial baroreceptors, that monitor the stretch exerted on the aortic arch^20^. Indeed, our follow-up *in situ* hybridization experiment revealed expression of PIEZO1/2 mRNAs within tdTomato-labeled nodose ganglionic neurons (**Fig. 1n-o**). In addition to PIEZO(s), transient receptor potential (TRP) cations channels have been implicated in baroreception^21^ and we observed that tdTomato-labeled neurons also expressed mRNAs encoding transient receptor potential ankyrin 1 (*Trpa1;* see **Fig. 1p**). These anatomical observations suggest that arterial baroreceptors express Oxtr(s), which conflicts with the results of Bai *et al* 2019 indicating that NDG^Oxtr^ specifically sense distention of the gastrointestinal tract^10^. It is possible that methodological differences, and in particular, the Cre-driver mice used to direct the expression of tdTomato to the *Oxtr* gene accounts for this discrepancy. Accordingly, we conducted an additional neuroanatomical experiment that delivered Cre-inducible AAV synthesizing tdTomato into the nodose ganglia of the Oxtr-Cre driver mice used by Bai et al 2019^10^. Using these mice, we again found that tdTomato clearly outlined a claw-like structure on the aortic arch (**Fig. 1q**). These anatomical results predict that NDG^Oxtr^ function as mechanoreceptors that monitor stretch exerted on the stomach, duodenum, and aortic arch.

### NDG^Oxtr^ that project to the gastrointestinal tract and aortic arch are anatomically segregated but each population likely expresses PIEZOs

Because tdTomato fibers originating from NDG^Oxtr^ were observed in the gastrointestinal tract and aortic arch, it is possible that both tissues are innervated by the same nodose ganglionic neurons. In other words, NDG^Oxtr^ innervating the gastrointestinal tract may also send collateral projections that target the aortic arch. To address this possibility, we delivered Oxtr^Cre^ mice retrograde Cre-inducible AAV synthesizing enhanced yellow fluorescent protein (eYFP) or tdTomato to the gastrointestinal tract or aortic arch, respectively (**Fig. 2a**). Three-weeks later, mice were perfused, and brain and nodose ganglia were extracted. As expected, a subset of soma in the nodose ganglia were labeled with eYFP or tdTomato; however, co-expression of the fluorophores was not observed (**Fig. 2b**). Confocal scans through thick coronal sections found that axons labeled with eYFP or tdTomato ascended through the rostral hindbrain (-7.0 mm from Bregma; **Fig. 2c**) and migrated dorsally and caudally toward the midline where they terminated within discrete regions of the caudal NTS (-7.5 to -7.8 mm from Bregma; **Fig 2c**); however, like the soma in nodose ganglia, colocalization of eYFP and tdTomato labeled fibers was not observed. These results are consistent with the notion that NDG^Oxtr^ send projections that innervate the stomach, duodenum, or aortic arch and suggest that separate populations of nodose ganglionic neurons target gastrointestinal or cardiovascular tissues.

**Fig. 2.**
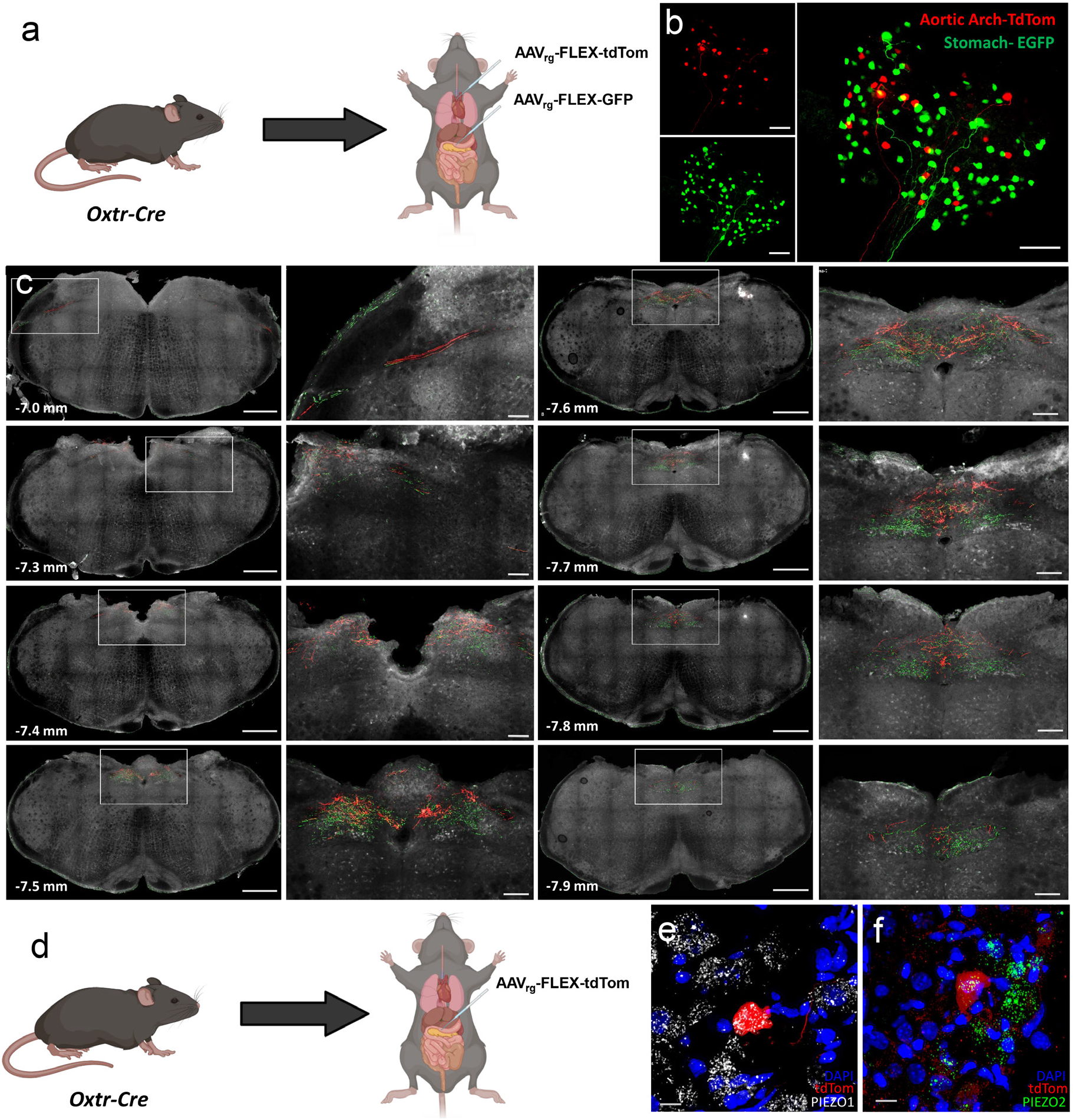
Separate populations of NDG^Oxtr^ innervate the stomach or aortic arch. (a) Schematic illustrating the application of AAV_rg_-Flex-tdTomato and -eGFP to the aortic arch and stomach, respectively (of Oxtr-Cre mice). (b) Confocal image of the whole NDG collected from one such mouse. (c) Images of the distribution of Aortic Arch-tdTomato and Stomach-eGFP fibers throughout the rostrocaudal extent of the NTS; rostrocaudal distance from bregma is denoted in the lower left-hand corner. (d) Schematic illustrating the application of AAV_rg_-Flex-tdTomato applied to the stomach. RNAscope *in situ* hybridization for (e) *Piezo1-* and (f) *Piezo2-* mRNA(s) in NDG^Oxtr^ that innervate the stomach (i.e., Stomach-tdTomato). Anatomical data are representative of 6 mice. Scale = 100 µm (b and c_insets_), 10 µm (e, f). Schematics made with Biorender.com.

As mentioned, nodose ganglionic neurons innervating the aortic arch utilize the mechanically gated ion channels, PIEZOs, to transduce vascular stretch into action potentials. Whether NDG^Oxtr^ similarly utilize PIEZO to monitor gastrointestinal distention is unknown. To begin to address this unknown, we delivered Oxtr^Cre^ male mice retrograde Cre-inducible AAV synthesizing tdTomato into the stomach and duodenum (**Fig. 2d**). Three-weeks later, mice were perfused and nodose ganglia were extracted and processed for *in situ* hybridization for *PIEZO1/2* mRNAs. **Figures 2e and 2f** depict confocal images of *PIEZO1* and *PIEZO2* mRNAs and nodose ganglionic neurons retrogradely labeled with tdTomato, and therefore, presumed to project to the gastrointestinal tract and synthesize Oxtr(s). Merged images reveal mRNAs coding for *PIEZO1* and *PIEZO2* within the confines of tdTomato labeled neurons [**Fig. 2e-f**]. Taken together, these results suggest that separate populations of NDG^Oxtr^ innervate the gastrointestinal tract and aortic arch; however, it is likely that subsets of each population utilize PIEZOs to monitor distention.

### Chemogenetic excitation of NDG^Oxtr^ decreases food intake and energy expenditure

The molecular and structural characteristics of NDG^Oxtr^ predict that these vagal sensory afferents monitor mechanosensation of the gut. To functionally test this prediction, we conducted experiments utilizing *in vivo* chemogenetics to selectively activate NDG^Oxtr^ while simultaneously recording metabolic parameters in conscious freely moving mice. Cre-inducible AAVs expressing DREADDs (Gq-DREADDs) and mCherry were bilaterally injected into the nodose ganglia of male Oxtr^Cre^ mice (**Fig. 3a**). This approach allows for selective chemogenetic excitation of NDG^Oxtr^ when CNO is administered. Male Oxtr^Cre^ mice given a Cre-inducible AAV only producing mCherry served as controls for the expression of Gq-DREADDs. Subsets of these male mice were also implanted with telemetry devices (DSI; HD-X10) and/or housed within a TSE PhenoMaster. After surgical recovery and acclimation to the TSE PhenoMaster, mice were given an injection of the saline vehicle or CNO (0.3 mg/kg, i.p.) at the onset of the dark phase.

**Fig. 3.**
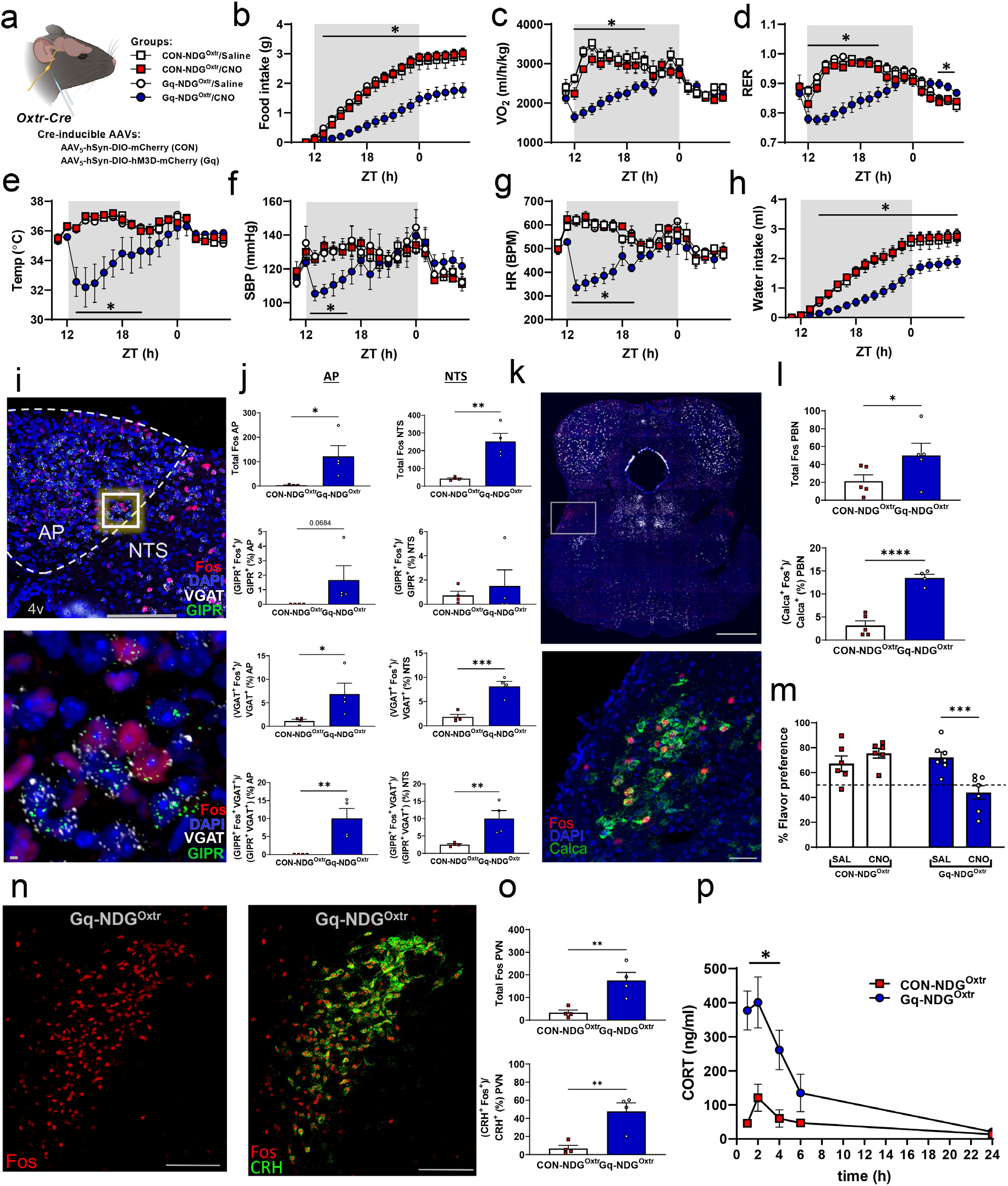
Acute stimulation of NDG^Oxtr^ affects cardiometabolic parameters and patterns of neuronal activation within brain regions implicated in conditioned taste avoidance and stress responses. **(a)** schematic of bilateral nodose injection treatment groups. **(b)** hourly food intake, **(c)** oxygen consumption (VO_2_) **(d)** respiratory exchange ratio (RER) **(e)** core body temperature, **(f)** systolic blood pressure **(g)** heart rate **(h)** hourly water intake **(i)** representative image of neuronal activation within the area postrema (AP) and nucleus of the solitary tract (NTS) 90 min following CNO administration **(j)** patterns of neuronal activation and colocalization of c-fos, GIPR, and VGAT within the AP (left panel) and NTS (right panel) following 90 min following CNO administration **(k)** representative image of neuronal activation and Calca colocalization within the parabrachial nucleus (PBN) 90 min following CNO administration **(l)** patterns of c-fos expression and colocalization with Calca-expressing neurons in the PBN 90 min following CNO administration **(m).** Change in flavor preference following conditioned taste avoidance paradigm **(n)** representative image of c-fos expression (left panel) and colocalization with CRH-expressing neurons in the paraventricular nucleus (PVN) of the hypothalamus 90 min following CNO administration **(o)** c-fos expression within the PVN (top panel) and colocalization of c-fos with CRH-expressing neurons within the PVN (bottom panel) **(p)** plasma corticosterone response to CNO (0.3 mg/kg i.p.) administration Data shown as mean ± s.e.m., n= 12 CON, 22 Gq **(b, c, d, h)**; n=4 CON, 5 Gq **(e, f, g)**; n= 6 CON, 7 Gq **(m)**; n= 5 CON, 5 Gq **(p)**. Statistical analysis performed by two-way analysis of variance (ANOVA) with Dunnett’s multiple comparisons test **(a, b, e, f, g, h)** or mixed effects analysis with Dunnett’s multiple comparisons testd **(c, d)**, two-way or repeated measures ANOVA with Fisher’s LSD **(p)**. Schematic **(a)** made with Biorender.com.

Filling of the gastrointestinal tract distends the muscular layers of the stomach and duodenum and this sensation is thought to contribute, at least in part, to meal termination and reduced food intake^22^. We hypothesized that chemogenetic excitation of NDG^Oxtr^ recapitulates the sensation of GI distention, and consistent with this hypothesis, administration of CNO to Oxtr^Cre^ mice given AAV-Gq-DREADDs significantly decreased food intake throughout the 18h testing period (**Fig. 3b**). Importantly, Oxtr^Cre^ mice given the AAV-mCherry control virus exhibited food intakes that were similar when administered CNO or the saline vehicle (**Fig. 3b**). These results suggest that chemogenetic excitation of NDG^Oxtr^ at the onset of the dark phase, when most of feeding occurs, potently suppresses food consumption, and it is unlikely that off target effects of CNO or Gq-DREAADs contribute to this effect.

We anticipated that mechanosensation of the gut would suppress feeding behavior but predicting how exciting NDG^Oxtr^ would affect energy expenditure was less clear. In rats, acute gastric filling is followed by post-prandial elevations in energy expenditure^23^; however, chronically enhancing gastrointestinal distention with inflation of a gastric balloon or bariatric surgery produces the opposite effect^24,25^. When compared to controls, Oxtr^Cre^ mice expressing Gq-DREADDs that were given CNO had significantly decreased, oxygen consumption (VO_2_; **Fig. 3c**), respiratory exchange ratio (RER; **Fig. 3d**), and core body temperature (**Fig. 3e**). These results suggest that exciting the NDG^Oxtr^ slows metabolic rate and is akin to manipulations that tonically enhance and/or simulate gastrointestinal distention. Moreover, the reduced RER observed shortly after delivery of CNO is indicative of the increased oxidation of fat that occurs during states of negative energy balance. Collectively, these results suggest that chemogenetic activation of NDG^Oxtr^ reduces indices of energy expenditure and promotes fat oxidation with the latter effect likely resulting from the reduced food intake that occurs shortly after Oxtr^Cre^ mice were administered CNO.

### Chemogenetic excitation of NDG^Oxtr^ decreases blood pressure, heart rate, and water intake

Elevated blood pressure activates arterial baroreceptors that provoke baroreflex-mediated reductions in cardiac output and vascular resistance. Subsets of NDG^Oxtr^ formed a claw-like structure on the aortic arch, suggesting that these sensory vagal afferents function as arterial baroreceptors. Here, we combine chemogenetics with telemetric recording of cardiovascular parameters to determine whether excitation of NDG^Oxtr^ mimics mechanical distention of the aortic arch and triggers baroreflex-mediated reductions in blood pressure and heart rate. As shown in **Figs. 3f-g**, 1 h after administration of CNO, Oxtr^Cre^ mice expressing Gq-DREADDs had significant and robust decreases in systolic blood pressure and heart rate relative to controls. At the nadir, systolic blood pressure and heart rate dropped ≈20 mmHg and ≈200 BPM, respectively. Remarkably, despite these robust effects, all mice fully recovered and were outwardly no different than controls after the effects of CNO had waned. In addition to perfusion pressure, arterial baroreceptors are thought to maintain intravascular blood volume by regulating fluid intake. Specifically, acute elevations in blood pressure or distending the vasculature via inflation of an atrial balloon significantly reduces water and saline consumption in rodents^26-28^. Similar to these prior studies, chemogenetic excitation of NDG^Oxtr^ significantly decreased water intake relative to controls (**Fig. 3h**). Taken together, these results indicate that exciting NDG^Oxtr^ elicits cardiovascular and behavioral responses that are reminiscent of those that follow loading of the arterial baroreceptors.

### Mechanosensation of the heart and gut engages neuronal circuits mediating satiety and stress responding

Next, we discerned the brain circuits that are engaged by chemogenetic excitation of NDG^Oxtr^. We again bilaterally delivered Cre-inducible AAV-GqDREADDs or AAV-mCherry control into the nodose ganglia of male Oxtr^Cre^ mice. After surgical recovery, mice were administered CNO (0.3 mg/kg, i.p.), and ninety-minutes later were euthanized, and brains were extracted and processed for dual Fos immunohistochemistry (IHC) and RNAscope *in situ* hybridization. As expected, giving CNO to mice that expressed Gq-DREADDs significantly increased Fos IHC in the area postrema (AP) and NTS relative to control mice expressing only mCherry (**Fig. 3i-j**). The AP and NTS mediate satiation^17,29^ and emerging work has shone a light on the glucose-dependent insulinotropic polypeptide receptor (GIPR) as a therapeutic target for obesity with agonism of this receptor or activation of GIPR-expressing neurons suppressing food intake^30,31^. Within the AP and NTS, GIPR(s) are primarily distributed on GABAergic neurons^32,33^, and fascinatingly, chemogenetic excitation of NDG^Oxtr^ significantly increased Fos IHC within cells synthesizing mRNAs encoding vesicular GABA transporter (*VGAT*; GABAergic marker) and *GIPR* (**Fig. 3i-j**). These results indicate that NDG^Oxtr^ demarcate vagal sensory afferents that excite neurons in the AP and NTS that synthesize GIPR(s) and GABA.

Neurons within the NTS send excitatory projections to the parabrachial nucleus (PBN), which relays sensory information to forebrain structures that shape the emotional valence of a stimulus^34^. Chemogenetic excitation of NDG^Oxtr^ augmented Fos IHC within the PBN (**Fig. 3k-l**), and intriguingly, we observed a significant increase in the number of neurons double-labeled for Fos IHC and mRNAs encoding calcitonin gene-related peptide (CGRP; *Calca* mRNA). The CGRP neurons of the PBN are activated by noxious stimuli^35^ and pairing their chemogenetic activation with consumption of a palatable sucrose solution elicits a conditioned taste avoidance^36^. These prior studies, in conjunction with the co-localization of Fos IHC and *Calca* mRNA, suggest that mechanosensation of the heart and gut may be a noxious stimulus. Towards this end, we paired chemogenetic excitation of NDG^Oxtr^ with consumption of a palatable saccharine solution to determine whether perceived mechanosensation of the heart and gut alters appetition of a hedonic stimulus. After consuming a palatable saccharin solution, Oxtr^Cre^ mice expressing Gq-DREAADD or mCherry were given the saline vehicle. As shown in **Fig. 3m** (left), 48 h later, both groups of mice exhibit a clear preference for a flavored saccharin solution over water during a two-bottle preference test. In contrast, 48 h after pairing saccharin with CNO, the preference for the flavored solution is completely abolished only in mice expressing Gq-DREADD (**Fig. 3m**; right). These results indicate that NDG^Oxtr^ recruit activation of *Calca* neurons residing within the PBN and suppress hedonically driven appetitive behavior.

The paraventricular nucleus of the hypothalamus (PVN) integrates interoceptive and exteroceptive signals and coordinates neuroendocrine responses that mitigate real or perceived threats to homeostasis. We found that chemogenetic activation of NDG^Oxtr^ robustly increased Fos IHC in the PVN (**Fig. 3n-o**). Dual IHC and RNAscope *in situ* hybridization revealed that chemogenetic excitation of NDG^Oxtr^ significantly increased Fos IHC within PVN neurons synthesizing mRNAs encoding corticotrophin-releasing-hormone (CRH; **Fig. 3n-o**). Within the PVN, neurons that synthesize CRH are known to activate the hypothalamic-pituitary-adrenal (HPA) axis, which elevates circulating levels of corticosterone (CORT). Corresponding experiments evaluated whether chemogenetic excitation of the NDG^Oxtr^ drives HPA axis activity to increase plasma levels of CORT. Gq-DREADDs or control mice were given CNO (0.3 mg/kg, i.p.) and blood was sampled, and plasma CORT was measured. Relative to controls, Oxtr^Cre^ mice expressing AAV-Gq-DREADD had robust and significant elevations in plasma corticosterone, 1 h after administration of CNO (**Fig. 3p**). To further characterize HPA axis activity, serial samples were collected 2, 4, 6 and 24 h after administration of CNO. Interestingly, relative to controls, Oxtr^Cre^ mice expressing AAV-Gq-DREADD had elevated plasma CORT 2, 4, and 6 h after administration of CNO with a return to basal levels occurring at 24 h (**Fig. 3p**). These results indicate that excitation of NDG^Oxtr^ is a stressor that potently activates the HPA axis and elevates plasma levels of CORT.

In mice, activation of CRH neurons in the PVN or CGRP neurons in the PBN is associated with vigilance that manifests as increased anxiety-like behavior^35,37,38^. To determine whether activation of NDG^Oxtr^ is anxiogenic, Gq-DREADDs or control mice were delivered CNO and 1 h later, anxiety-like behavior was assessed using the light-dark box (LDB), elevated plus maze (EPM), and open field arena (OFA). As depicted in **Supp. Fig. 1**, Oxtr^Cre^ mice expressing AAV-Gq-DREADD that were given CNO had fewer entries in the light- and spent less time on the light-side of the LDB when compared to that of controls. Chemogenetic excitation of NDG^Oxtr^ also reduced entries into the open arms or center area of the EPM or OFA, respectively. The total distance travelled was also significantly reduced by chemogenetic activation of NDG^Oxtr^. Collectively, these results suggest driving NDG^Oxtr^ engages brain circuits that suppress feeding, alter the valence of hedonic stimuli, elevate circulating stress hormones, and promote behavioral vigilance.

### Repeated chemogenetic excitation of NDG^Oxtr^ decreases food intake and body mass but does not affect anxiety-like behavior

The hypometabolism and vigilance that we observed during excitation of NDG^Oxtr^ closely resembles torpor, which is a state that heterotherms, like mice, enter when the availability of metabolic fuel is insufficient to meet real or perceived energetic demands^12^. Because torpor slows metabolic rate and alters fuel utilization, methods that artificially induce torpor have been explored for a variety of applications^39-41^. In this regard, the reductions in food intake, heart rate and energy expenditure that accompany excitation of NDG^Oxtr^ present the possibility that these sensory vagal afferents may be repeatedly engaged to artificially induced torpor and long-term changes in energy balance. To address this possibility, we again delivered Cre-inducible AAV expressing Gq-DREADDs and mCherry bilaterally into the nodose ganglia of male Oxtr^Cre^ mice. Littermates that only expressed mCherry served as controls. Following surgical recovery, mice were implanted with telemetry devices to enable continuous recording of cardiovascular parameters. After habituation to the TSE PhenoMaster, mice were delivered CNO (0.3 mg/kg, i.p.) at the onset of the dark phase (11:00 h) every-other-day for eleven days and food intake, body weight, indices of energy expenditure, and cardiovascular parameters were recorded. As shown in **Fig. 4**, on days when CNO was given, mice expressing AAV-Gq-DREADDs had decreased food intake (**Fig. 4a**), body temperature (**Fig. 4b**), VO_2_ (**Fig. 4e**), and RER (**Fig. 4f**) when compared to baseline and non-injection days. Importantly, these alterations were not observed in control mice expressing AAV-Gq-mCherry (**Supp. Fig. 2**). That is, food intake, body temperature, VO_2_, and RER were not altered subsequent to CNO administration in these control mice. These results suggest that the effects of CNO do not linger and the prior reduction in food intake observed during chemogenetic excitation of NDG^Oxtr^ does not elicit compensatory overconsumption of calories or changes in energy expenditure. Regarding cardiovascular parameters, on days when CNO was given, mice expressing AAV-Gq-DREADDs had decreased blood pressure (**Fig. 4i**) and heart rate (**Fig. 4j**) when compared to non-injection days or baseline measures. However, blood pressure, and heart rate were similar amongst the injection, non-injection and baseline days in the control mice. Intriguingly, repeated chemogenetic excitation of NDG^Oxtr^ produced a 3-5% reduction in body mass that was sustained throughout the experimental paradigm **(Fig. 4m**). Collectively, these results indicate that repeatedly exciting NDG^Oxtr^ promotes sustained decreases in body mass but chronic reductions in energy expenditure and cardiovascular parameters were not observed.

**Fig. 4.**
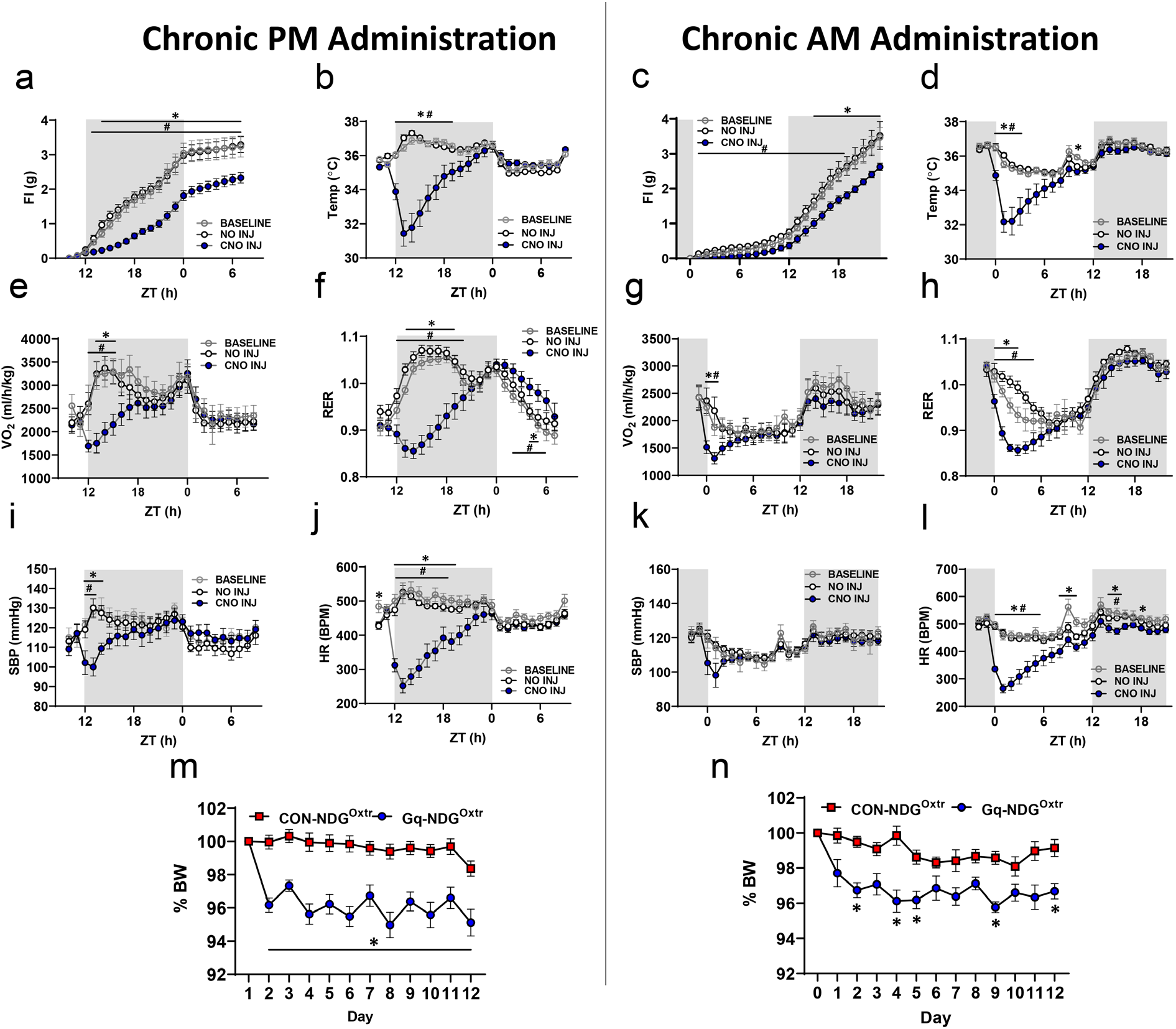
Chronic activation of NDG^Oxtr^ affects cardiometabolic parameters, resulting in persistent reductions in body weight. CNO (0.3 mg/kg, i.p.) was administered every other day for 11 days at ZT 11 (1h before lights off, left panel) or ZT23 (1h before lights on, right panel). Chronic administration reduced food intake **(a,c),** temperature **(b,d)** metabolic **(e-h)** and cardiovascular parameters **(i-l)**, and body weight **(m,n)**. Gq-NDG^Oxtr^ data are averaged between baseline (2 days), no injection (5 days) and CNO injection days (6 days). Data shown as mean ± s.e.m., **(a,e,f)** n= 13 Gq; **(b,I,j)** n=8 Gq **(m)** n=17 CON, 13 Gq **(n)** n=11 CON, 8 Gq. 2 way RM ANOVA or mixed effects model with Dunnett’s multiple comparisons tests **(a-l)** or mixed effects model with Šídák’s multiple comparisons test **(m,n)**. *, p<0.05 between BASELINE and CNO INJ; #, p<0.05 between NO INJ and CNO INJ.

In mice, chronic unpredictable stress is accompanied by decreased body weight, increased anxiety-like behavior, and augmented HPA axis activity^42,43^. During chemogenetic excitation of NDG^Oxtr^, mice display an anxiogenic phenotype and greatly elevated plasma CORT, a proxy of HPA axis activity. Therefore, it is possible that the repeated excitation of NDG^Oxtr^ that promotes weight loss is perceived as chronic unpredictable stress. To address this possibility, after the 7^th^ day, on the even numbered days when CNO was not given (i.e., days 8, 10 and 12), mice were tested for anxiety-like behavior to determine whether a history of chemogenetic excitation of NDG^Oxtr^ alters mood and affect. As depicted in **Supp. Fig. 3a-f**, control mice and those given AAV-Gq-DREADDs behaved similarly in the LDB and EPM on even numbered days when CNO was not administered. Next, we considered that oxytocin is a potent mediator of social behavior^44^ and we used the social interaction paradigm to assess indices of social avoidance or social fear^45^. Controls and Gq-DREADDs mice that were previously treated with CNO spent similar amounts of time investigating novel conspecifics, suggesting that repeated chemogenetic excitation of NDG^Oxtr^ had no effect on social behavior (**Supp. Fig. 3g-i**). Taken together, these results demonstrate that repeated chemogenetic excitation of NDG^Oxtr^ alters food intake and energy expenditure to reduce body mass; however, these episodes of NDG^Oxtr^ activation are not associated with heightened stress responsivity as measured by anxiety-like behaviors.

### The reduced food intake and body mass that accompanies repeated chemogenetic activation of NDG^Oxtr^ does not require temporally coupling torpor with eating behavior

Finally, we contemplated that torpor shares several commonalities with sleep and when faced with energetic challenges, mice will tend to enter torpor at the onset of the light-phase, during their circadian nadir of activity^46^. Taking this into account, we evaluated whether repeatedly delivering CNO at the onset of the light-phase would recapitulate the cardiometabolic changes that were previously observed. Mice were delivered AAV-Gq-DREADDs or AAV-mCherry, implanted with telemetry devices and housed in the TSE Phenomaster as described. Following surgical recovery and habituation to the TSE PhenoMaster, mice were delivered CNO (0.3 mg/kg i.p.) at the onset of the light phase (ZT 23h), every-other-day for 11 days. As with our previous experiments, administration of CNO to mice expressing AAV-Gq-DREADDs decreased food intake (**Fig. 4c**), body temperature (**Fig. 4d**), VO_2_ (**Fig. 4g**), RER (**Fig. 4h**), blood pressure (**Fig. 4k**) and heart rate (**Fig. 4l**) when compared to days that CNO wasn’t injected and to baseline measurements; however, many of these effects were now constrained to the light-phase. These alterations were not observed in controls (**Supp. Fig. 3**). Remarkably, on the days that CNO was delivered, chemogenetic excitation of NDG^Oxtr^ significantly suppressed food intake and this effect persisted throughout the dark-phase when measures of cardiovascular function were largely similar amongst the conditions (**Fig. 4**). Similar to results observed during the Chronic PM administration of CNO (**Fig. 4m**), alterations in energy balance during Chronic AM CNO administration led to a 3-4% reduction in body mass (**Fig. 4n**). Collectively, these results indicate that the reductions in food intake that follow excitation of NDG^Oxtr^ cannot be attributed to decreased blood pressure and heart rate. Moreover, chemogenetic excitation of NDG^Oxtr^ does not need to be concurrent with ingestive behavior in order to observe suppressed food intake. Rather, the excitation of NDG^Oxtr^ may engage brain circuits that mediate the long-term regulation of energy balance.

## DISCUSSION

Here, we reveal that Oxtr(s) are expressed on nodose ganglionic neurons that monitor mechanosensation of the heart and gut. Our anatomical experiments found that NDG^Oxtr^ comprise a subset (≈20%) of nodose ganglionic neurons innervating the aortic arch or gastrointestinal tract that express molecular markers and structural endings indicative of mechanosensation. Fascinatingly, chemogenetic excitation of NDG^Oxtr^ produced a torpor-like phenotype that includes suppression of eating and drinking as well as robust decreases in blood pressure, heart rate, body temperature, and energy expenditure. Activation of NDG^Oxtr^ was coupled to augmented Fos expression within brain circuits mediating stress responding and this expression was predictive of conditioned taste avoidance, augmented HPA axis activity, and increased anxiety-like behavior. Repeatedly coupling excitation of NDG^Oxtr^ with ingestive behavior by delivering CNO at the onset of the dark-phase produced sustained reductions in body mass without affecting anxiety-like behavior. Remarkably, delivering CNO at the onset of the light-phase, when mice are inactive or sleeping, suppressed food intake during the subsequent dark-phase and this also produced sustained reductions in body mass. Collectively, these results demonstrate that vagal afferents transducing mechanosensation of the heart and gut have a profound influence over brain circuits mediating cardiometabolic function and stress responding. The broad implication is that NDG^Oxtr^ govern interoceptive pathways that can be leveraged to understand and alleviate cardiometabolic diseases and mental health disorders.

The brain monitors the moment-to-moment status of blood volume and perfusion pressure, in part, via arterial baroreceptors whose soma reside within the NDG^47^. Arterial baroreceptors form a claw-like structure that encapsulates the aortic arch and transduces stretch exerted on the vessel wall into action potentials that are carried to the NTS^20,48,49^. While still an active area of research^21,50^, recent evidence points to mechanosensitive ion channels, PIEZO(s), as molecular markers for arterial baroreceptors^51^. We found that NDG^Oxtr^ express *PIEZO1/2* mRNAs and send projections to the aortic arch that form a claw-like structure. Chemogenetic excitation of NDG^Oxtr^ significantly reduced blood pressure and heart rate, which is reminiscent of baroreflex activation elicited by hypertensive stimuli. We also found that chemogenetic excitation of NDG^Oxtr^ suppressed water intake. This presents somewhat of a conundrum because the unloading of arterial baroreceptors during hypotension drives^52^, but the loading that occurs with hypertension abrogates, fluid intake^53^. Thus, the suppressed water intake that we observed may appear paradoxical because chemogenetic excitation of NDG^Oxtr^ produced hypotension; however, our interpretation of these results is that activation of NDG^Oxtr^ recapitulates the sensation of baroreceptor loading and this blunts thirst and drinking behavior.

Studies using electrophysiological recordings found that distention of the GI tract elicited firing of vagal afferents that persisted after ablation of the mucosal surface^54-57^. This suggested that vagal mechanoreceptors were not located in the mucosa, but rather, resided within the muscle wall of the gastrointestinal tract. Indeed, anatomical studies revealed the presence of IGLE(s) and IMA(s) within the circular and longitudinal muscle layers^16,58^; though, the strongest evidence linking IGLE(s) to gastrointestinal mechanosensation comes from experiments that found that stretch sensitive vagal afferents were associated with receptive fields on the stomach that were innervated by IGLE(s)^19,59,60^. Consistent with the results of Bai et al.^10^, we report that NDG^Oxtr^ give rise to IGLE(s) and IMA(s) that populate the stomach and densely occupy portions of the duodenum that are proximal to the pylorus. Additionally, NDG^Oxtr^ innervating the gastrointestinal tract synthesize *PIEZO1/2* mRNAs, which were recently found to transduce stomach distention and mediate feeding in *Drosophila*^61,62^. Our results, in conjunction with those from prior electrophysiological and anatomical studies, support the notion that Oxtr(s) demarcate vagal afferents subserving mechanosensation of the gastrointestinal tract. Interestingly, we found that, relative to males, IGLE(s) originating from NDG^Oxtr^ were less dense in females. Peripherally administered oxytocin appears to be more efficacious at reducing food intake in males relative to females^63,64^ and the scant number of IGLE(s) that we observed in the proximal duodenum of females may contribute to this effect.

Vagal sensory afferents form excitatory synapses onto cells within the NTS^48^ and these cells, deemed 2^nd^ order neurons, relay interoceptive signals to nuclei orchestrating body-to-brain communication. Prior work from our group revealed that NDG^Oxtr^ synapse onto 2^nd^ order neurons that synthesize preproglucagon (PPG)^65^. These PPG neurons are thought to release glucagon-like peptide 1 (GLP-1), which stimulates glucagon-like peptide 1 receptors (GLP-1R) expressed throughout the brain^66^. Relevant to the current study, peripherally administered oxytocin suppresses feeding by acting on vagal afferents^14^ and PPG neurons are required for this anorexigenic effect^65^. Interestingly, PPG neurons are activated by gastric distention as well as psychogenic stressors. Moreover, GLP-1 labeled fibers and GLP-1R(s) are distributed throughout brain regions governing physiological and behavioral response to stress^66^. Central administration of GLP1-R agonists lower core body temperature^67^, which is similar to the reduction in core body temperature that we observed when NDG^Oxtr^ are excited. Therefore, it is possible that the anorexigenic and stress responsive phenotype that accompanies chemogenetic excitation of NDG^Oxtr^ can be attributed, in part, to the engagement of PPG neurons. However, the bradycardia that we observed is contrary to what occurs with engagement of the central GLP-1 system^68^, and therefore, it is likely that NDG^Oxtr^ recruit diverse populations of 2^nd^ order neurons to produce the complex phenotype that follows their excitation.

The hypometabolism and vigilance that we observed during excitation of NDG^Oxtr^ closely resembles torpor, which is a state that heterotherms, like mice, enter when the availability of metabolic fuel is insufficient to meet real or perceived energetic demands. Reductions in body temperature^69^, reduced cardiac output and perfusion pressure^70^, decreased respiratory rate and oxygen consumption^71,72^, suppression of hunger^73^ or thirst^74^, and behavioral vigilance^73^ have all been reported as torpor-induced effects. Excitation of NDG^Oxtr^ similarly produced these effects and, like torpor^75,76^, our effects persisted for 4-6 hours. Food scarcity, cold and/or threat of predation are conditions that promote torpor, and by convention, nervous system connections that sense the external environment are studied as mediators. Here, we demonstrate that exciting vagal afferents relaying mechanosensation of the heart and gut potently elicits a torpor-like phenotype. Recently, Matsuo and colleagues reported that vagal afferents partly contribute to the torpor-like phenotype that accompanies systemic delivery of Trpa1 agonists^71^, and intriguingly, we determined that ≈30% of NDG^Oxtr^ expressed *Trpa1* mRNA. Taken together, these results indicate that excitation of NDG^Oxtr^ is sufficient to drive physiological and behavioral responses that are hallmarks of torpor and further suggest that interoceptive pathways potently alter cardiometabolic function and behavior in response to real or perceived threats to the internal milieu.

Artificially-induced torpor has been proposed for applications ranging from medical therapies to space travel^39-41^. These applications often require prolonged bouts of torpor, and as proof of concept, we evaluated the effects that repeated activation of NDG^Oxtr^ had on cardiometabolic function and stress responding. Artificially inducing torpor by repeatedly exciting NDG^Oxtr^ promoted sustained weight loss but transiently lowered energy expenditure, body temperature and heart rate. These results suggest the weight loss can likely be attributed to the blunted food intake that occurs on the day of NDG^Oxtr^ excitation. The repeated bouts of ‘artificial torpor’ were not associated with exaggerated indices of stress responding (e.g., anxiety-like behavior), suggesting that the intervention is well-tolerated. Intriguingly, weight loss was also observed when the activation of NDG^Oxtr^ and the circadian rhythm of ingestive behavior were temporally uncoupled. That is, exciting NDG^Oxtr^ at the onset of the light-phase, when mice are inactive or sleeping but not eating, promoted sustained weight loss. Decreased food intake also contributed to this weight loss; however, this anorectic effect occurred 12 hours after CNO administration when the hypometabolism that follows excitation of NDG^Oxtr^ had subsided. Taken together, these results establish that repeated activation of NDG^Oxtr^ is well-tolerated and promotes decreased food intake that contributes to sustained reductions in body mass. The implication is that NDG^Oxtr^ may be a promising therapeutic target for the treatment of cardiometabolic disorders, like obesity. However, it should be noted that the suppressed feeding behavior and weight loss was observed in lean mice maintained on a standard diet. It is possible that artificially inducing torpor with activation of NDG^Oxtr^ may produce larger reductions in body mass when mice are rendered obese on a high-fat diet. Alternatively, maintenance on a palatable diet may override the mechanically gated satiation signals supplied by NDG^Oxtr^ but still engage mechanosensitive pathways that promote hypometabolism and vigilance. In this way, overconsumption of palatable and calorically dense foods and fluids may promote states of positive energy balance and affective disorders, like obesity and anxiety, by simultaneously overwhelming satiety-signals supplied by mechanosensation of the heart and gut while also recruiting neuronal pathways promoting energy conservation and stress responding.

## METHODS

### Animals

All procedures were approved by the Institutional Animal Care and Use Committee at the University of Florida. Mice were on a C57BL/6J background, housed individually, and maintained on a 12:12 h light/dark cycle, in temperature (20-26°C) and humidity (30-70%)-controlled rooms. Additionally, subjects were given *ad libitum* access to standard rodent chow (Envigo Teklad [7912]) and water, unless stated otherwise.

The majority of the studies were performed using *Oxtr*-Cre mice (Mutant Mouse Resource Research Centers, Stock #036545-UCD). This bacterial artificial chromosome transgenic mouse line expresses Cre recombinase under control of the *Oxtr*-specific promoter and has been described previously^77,78^.

In addition, a cohort of Oxtr-T2A-Cre-D (Jax Strain #031303) was utilized for anatomical tracing studies^79^.

### Adenoassociated viruses (AAVs)

Several Cre-inducible AAVs were used to allow for interrogating the characteristics of *Oxtr*-expressing neurons of the NDG (NDG^Oxtr^). These AAVs were obtained from Addgene and were delivered into the NDG (or to peripheral tissues innervated by NDG) of *Oxtr*-Cre mice. Specifically, AAV_PHP.S_-FLEX-tdTomato (Addgene, Catalog #28306-PHP.S) was injected directly into the NDG and was used for anterograde tracing studies. AAV_2/retro_-FLEX-tdTomato (Addgene, Catalog #28306-AAVrg) and AAV_2/retro_-pCAG-FLEX-EGFP-WPRE (Addgene, Catalog #51502-AAVrg) were used as retrograde tracers to evaluate NDG^Oxtr^ innervation of the GI tract and the aortic arch. Finally, pAAV_5_-hSyn-DIO-hM3D(Gq)-mCherry (Catalog #44361-AAV5) and pAAV_5_-hSyn-DIO-mCherry (Catalog #50459-AAV5) were injected directly into the NDG in order to perform chemogenetic studies that evaluate the functionality of NDG^Oxtr^.

### Surgical procedures

#### Pre- and post-operative procedures

For all survival surgeries, mice were anesthetized using 2% isoflurane and then administered analgesic (Buprenorphine SR-LAB (Zoo Pharm), 1 mg/kg, s.c.; meloxicam (Pivetal Alloxate), 20 mg/kg, s.c.). Body temperatures were then maintained on a heating pad throughout the duration of the surgery and ophthalmic ointment was applied for corneal protection. Finally, surgical site(s) were prepared for incision by removing the fur and disinfecting skin. Post-operative care: in addition to pre-operative analgesia, mice received additional doses of buprenorphine SR-LAB 48 h post-procedure, and meloxicam 24, 48, and 72 h post-procedure.

#### AAV delivery to the NDG

After performing pre-operative procedures, *Oxtr-Cre* mice were placed in dorsal recumbency, and a longitudinal incision was made to expose the salivary glands. The vagus nerve was separated from the carotid artery until the nodose ganglion became accessible. Cre-dependent AAVs [AAV_PHP.S_-FLEX-tdTomato (Addgene, Catalog #28306-PHP.S), pAAV_5_-hSyn-DIO-hM3D(Gq)-mCherry (Catalog #44361-AAV5) or pAAV_5_-hSyn-DIO-mCherry (Catalog #50459-AAV5)] were pulled into beveled glass pipettes (Sutter Instrument, O.D.: 1.0 mm, I.D.: 0.58 mm) and injected (∼0.3-0.5 µL/site) into NDG on each side via pressure injection system (PV830 Pneumatic PicoPump, WPI). The incision site was sutured and mice were returned to home cages for recovery.

#### AAV delivery to the Aortic Arch

After performing pre-operative procedures, a subset of *Oxtr-Cre* mice was placed in dorsal recumbency and then delivered AAV_2/retro_-FLEX-tdTomato (Addgene, Catalog #28306-AAVrg), onto the aortic arch following surgical procedures previously described^5^. Briefly, following intubation for artificial ventilation (tidal vol; 0.2 ml, min vol; 26 ml min^-1^, Pressure 21 cmH_2_O), a longitudinal midline cervical incision was made from the sternal notch to mid-chest and pre-trachea muscles bluntly separated. A sternotomy was then performed (approximately 3-4 mm) to expose the aorta and pericardial cavity. A pulled glass micropipet was then used to apply 1 ul of virus to the aortic arch. Viral constructs were left *in situ* for 5 minutes to allow for the diffusion of AAVrg into the aortic arch. The incision site was then sutured closed, and the mouse either returned to a recovery cage or then delivered virus to the GI tract.

#### AAV delivery to the GI tract

After performing pre-operative procedures, a longitudinal laparotomy was performed and the stomach and proximal duodenum exposed and isolated. Cre-dependent AAV [5 µl; AAV_2/retro_-FLEX-tdTomato (Addgene, Catalog #28306-AAVrg) or AAV_2/retro_-pCAG-FLEX-EGFP-WPRE (Addgene, Catalog #51502-AAVrg)] was then applied to the surface of the antrum of the stomach, the pylorus and the proximal duodenum using a disposable pipet. The incisions were sutured and mice were returned to home cages subsequently for recovery.

#### Telemetry device implantation

In order to evaluate cardiovascular parameters during chemogenetic excitation of NDG^Oxtr^, a subset of mice (n = 24) that were previously administered AAVs into the NDG, underwent a further surgery to implant radiotelemetry devices (HDX10; Data Sciences International, St. Paul, MN). These surgeries were performed as previously-described^37^, using the anesthesia and analgesia procedures that are described above. Briefly, radiotelemetry transmitters were placed intraperitoneally and secured to the abdominal wall using suture. The abdominal muscles were then closed with absorbable suture. Using the same procedure as for insertion of the Millar catheter, the fluid-filled catheter was inserted into the distal left carotid artery and secured in place with suture. The skin was closed with non-absorbable 5-0 monofilament suture.

### Indirect calorimetry and ingestive behavior

Three weeks after surgical procedures, mice were transferred into an automated indirect calorimetric system (PhenoMaster, TSE systems) that recorded oxygen consumption (VO_2_), respiratory exchange ratio (RER) and continuous food/water intake. RER is the ratio of carbon dioxide production to oxygen consumption (VCO_2_ / VO_2_). The environmental temperature (22-25℃), light (12:12 h light/dark cycle) and humidity (50%) were maintained by way of the climate-control chamber and TSE PhenoMaster software. Mice were habituated in the chamber for at least three days prior to any experimental data collection.

### Telemetric cardiovascular recording

During cardiovascular recordings, mice were also housed in the TSE PhenoMaster (as described above), with their cages placed on PhysioTel® receivers (Data Sciences International, St. Paul, MN). For the duration of the cardiovascular studies, systolic blood pressure (SBP), diastolic blood pressure (DBP), mean arterial pressure (MAP), heart rate (HR) and core body temperature were continuously acquired and hourly bin data were calculated using Ponemah Software (Data Sciences International, St. Paul, MN).

### Acute CNO administration

For acute experiments, food access was restricted for 2 h prior to dark phase. After at least 3 days habituation to i.p. injections (0.1 ml saline), mice received one injection of 0.1 ml CNO (0.3 mg/kg, i.p.) 1h before onset of the dark phase (zeitgeber time, ZT, 11). Access to food was given at the onset of the dark phase, and food intake, VO_2_ and RER recorded. A subset of mice (n= 4 CON-NDG^Oxtr^ and 5 Gq-NDG^Oxtr^) had telemetry devices implanted for assessment of cardiovascular parameters.

### Chronic CNO administration

Mice were given at least 3 days to habituate to the Phenomaster cages and saline injections (0.1 ml i.p.). CNO (0.3 mg/kg i.p.) was injected every other day for 11 days (6 CNO injections), 1h before onset of the dark phase (ZT 11, for chronic PM CNO administration) or 1h before onset of the light phase (ZT 23, for chronic AM CNO administration). VO2, RER and food intakes were measured during this time. A subset of these mice (n=15; 7 CON-NDG^Oxtr^ and 8 Gq-NDG^Oxtr^) had telemetry implants in order to record cardiovascular parameters.

### Conditioned taste avoidance

To determine whether acute activation of NDG^Oxtr^ was aversive, a conditioned taste avoidance (CTA) test was performed using a 2-bottle intake chamber (52 cm x 10 cm x 30.5 cm, l x w x h) with lickometry devices (Noldus) using a protocol adapted from previously published CTA experiments^80,81^. Prior to the experiment, mice were habituated to the intake chamber for 1h per day for 3 consecutive days with *ad libitum* access to a water bottle. Mice were not water deprived during this habituation phase. This was then followed by a training period; in order to motivate the mice to drink from the bottles during a short period of time, mice were water restricted overnight (<16h) every other night, prior to the intake session. During the training session, a water bottle was placed in one of the 2 bottle ports and mice were given 30 min to drink. The location of the bottle was alternated and counterbalanced between sessions to limit side preferences. Mice were then returned to their home cage. Two hours following the end of the session, food and water were returned. This habituation period took 4-7 days; conditioning did not proceed until water intakes stabilized over the 30 min session. For conditioning, 2 different palatable solutions of 0.15% saccharine flavored with 0.05% Kool-Aid were used. On the first day of conditioning (following overnight water deprivation), the mice were given 30 min to consume flavored solution 1, and then injected with vehicle (saline) and returned to the home cage. The 2nd day was a rest day, and the mouse was then water deprived overnight and tested for preference on the 3^rd^ day, where the mouse was given 30 min access to both a water bottle and a bottle containing flavor 1. Mice were given another rest day, and then underwent conditioning to flavor 2 in the same manner as flavor 1; however, the 2^nd^ flavor was paired with CNO (0.3 mg/kg), rather than saline. They were given a day off after flavor 2 conditioning, and were tested on the following day with access to bottles containing water and flavor 2. Flavors 1 and 2 were counterbalanced to account for any differences in palatability. Preference was calculated as licks for flavored solution divided by the total licks for water and flavored solution combined.

### Open Field Test

The open field (OF) test was conducted in a 45 x 45 x 30 cm (l x w x h) square arena constructed of white plastic floor and walls. Each mouse was brought into the procedure room immediately before the test and placed in the center of the open field arena 30 min after i.p. CNO (0.3 mg/kg) injection. Results including total distance travelled and entries into center (located in the center of arena with 50% of total area) or edge were recorded and analyzed using Ethovision XT 13 software (Noldus Information Technology, Netherlands). The total duration of open field test was 5 min.

### Elevated Plus Maze Test-Acute and chronic activation

The elevated plus maze (EPM) arena has two opposing open arms and two opposing closed arms, raised 61 cm above the floor. All arms are 35 x 5 cm (l x w) with white floors, and the walls of the closed arms are made of black polycarbonate, 20 cm high. The test started 2 h after the end of the dark cycle. The total duration of the test was 5 min. Results, including the number of open arm entries and distance travelled in open and closed arms (excluding center) were recorded and analyzed by Ethovision XT 13 (Noldus Information Technology, Netherlands). Acute administration: 30 min after i.p. CNO (0.3 mg/kg) injection, the test mouse was placed in the center of the maze and allowed free exploration of the arena. Chronic administration: the experimental mouse was run through the EPM on the morning following PM CNO (0.3 mg/kg, i.p.) administration (approximately 14 h after administration). EPM was performed on Day 8 of chronic CNO treatment.

### Light Dark Box Test-Acute and chronic activation

Approximately 2h after the start of the light phase, mice were placed in a light dark box (LDB) test arena in which one chamber (20 x 40 x 35 cm, l x w x h) is kept dark, with black polycarbonate walls and lid, and the light side (20 x 40 x 35 cm) is made of clear polycarbonate walls with no lid; the floor of both sides is grey. The two chambers are separated by a black wall with a small central opening (7 x 7 cm), which the test mouse could freely cross through. Mice were initially placed into the middle of the dark chamber, facing the back wall, and allowed 5 min of free exploration. Tests were digitally-recorded and hand-scored. Acute administration: 30 min after i.p. CNO (0.3 mg/kg) injection, the test mouse was placed in the center of the dark chamber and allowed free exploration of the arena. Chronic administration: the experimental mouse was tested in the LDB on the morning following PM CNO (0.3 mg/kg, i.p.) administration (approximately 14h after administration). LDB was performed on day 10 of chronic CNO treatment.

### Three-chambered Social Interaction Test

To determine whether chronic activation of NDG^Oxtr^ affected social behavior, a 3-chamber social interaction test (3CSIT) was run on D12 of chronic stimulation. The polycarbonate arena (60 cm x 40.5 cm x 22cm, l x w x h) is divided into 3-19 cm wide chambers with doors between them. During the first phase of the test, the mouse was given 5 min to investigate the empty arena. For the social preference test, the mouse was returned to the center chamber and a clear acrylic cage (10 cm diameter, 20 cm tall) was placed in each end chamber. One chamber had an empty cage, and a novel C57BL6/J mouse was placed in the other cage. The experimental mouse was then allowed to explore the chambers for 5 minutes. Both phases were recorded and scored using Ethovision XT13 (Noldus, the Netherlands). Time spent in proximity to the cage was measured, and social preference was calculated as time spent near the mouse divided by total time spent near the mouse and near the empty cage. The test was performed 2 h after onset of the light cycle, approximately 14 h after the final (D11) PM administration of CNO.

### Assessment of Plasma CORT

One hour after i.p. CNO (0.3 mg/kg) injection, blood samples from the tail tip were collected from CON-NDG^Oxtr^ and Gq-NDG^Oxtr^ mice; additional samples were collected 2, 4, 6, and 24 h following injection. Blood samples were collected into EDTA-treated tubes and centrifuged at 2500 x G for 15 min at 4°C, and isolated plasma samples stored at -80°C. Plasma corticosterone (CORT) levels were measured using an ^125^I radioimmunoassay kit (MP Biomedicals, Orangeburg, NY) as previously described^82^.

### Tissue collection and sectioning

At the end of the studies, mice were injected with 390 mg sodium pentobarbital/50 mg phenytoin per ml solution (Euthanasia-III Solution, Med-Pharmex Inc., Pomona, CA), 0.1 ml i.p.). Once breathing ceased, transcardial perfusions were performed by clearing with 0.15M NaCl followed by fixation with 4% paraformaldehyde (Sigma Aldrich). Brains were extracted and post-fixed in 4% paraformaldehyde for 4 h. After post-fixation, brains were transferred into 30% sucrose and stored at 4°C, until sectioned for either *in situ* hybridization or immunohistochemistry. The left and right NDG were dissected and placed in in 4% paraformaldehyde for 5 min before they were stored in 20% RNase-free sucrose at 4 °C, until being whole-mount imaged before being sectioned for *in situ* hybridization. The aortic arch and stomach were also carefully dissected, collected and stored in RNase-free saline at 4 °C, until being whole-mount imaged.

Brains and NDG for *in situ* hybridization were sectioned at a thickness of 20 µm or 10 µm respectively, using a Leica cryostat (Leica CM3050S, Leica). The cryostat and other tools were cleaned with RNase remover (Fisher Scientific) and ethanol before use. Tissues were first collected in RNase-free PBS and then mounted on slides (Tissue Path Superfrost Plus Gold Microscope Slides, Fisher Scientific). After air-drying at room temperature for 1 h, slides were rinsed with PBS, incubated in 4% PFA for 15 min, and then dehydrated for 5 min in 50%, 75%, and 100% EtOH at room temperature. Following that, slides were incubated in H_2_O_2_ for 10 min, rinsed with PBS, and allowed to dry for 10-15 min before being stored at - 80°C for further processing. Brains for immunohistochemistry experiments were sectioned at 30 µm and stored in cryoprotective solution (1L 0.1M PBS, 20g PVP-40, 600ml ethylene glycol, 600g sucrose) at -20°C.

### *In situ* hybridization

*In situ* hybridization with various probes was performed to examine Cre recombinase expression in *Oxtr*-Cre mice and the phenotype of *Oxtr*-expressing neurons using the RNAscope^®^ V2 Multiplex Fluorescent Reagent Kit (Advanced Cell Diagnostics, Newark, CA). Detailed procedures and the reagents are provided as per the manufacturer’s instructions (Advanced Cell Diagnostics, Inc., Hayward, CA) and as previously described ^83^. Briefly, brain and NDG slides were first treated with protease IV or III respectively (Advanced Cell Diagnostics, Lot#2002337) for 20 min in the dark at room temperature. After 3 consecutive washes with RNase-free water, brain slides were incubated in probes for the mRNA(s) of vGat (Mm-Slc32a1, Cat. No. 319191), GIPR (Mm-Gipr, Cat. No. 319121), Calca (Mm-Calca, Cat. No. 578771), CRH (Mm-Crh, Cat. No. 316091), whereas NDG slides were incubated in probes for the mRNA(s) of Oxtr (Mm-oxtr, Cat. No. 412171), PIEZO-1 (Mm-Piezo1-O2; Cat. No.529091), and PIEZO-2 (Mm-Piezo2-E39-E44; Cat. N. 433421), TRPA1 (Mm-Trpa1-C2; Cat. No.400211-C2) (working dilution 1:50, Advanced Cell Diagnostics) in probe diluent for 2 hours at 40°C. The probes contained double Z shaped probe pairs, which are complementary to target RNA, and thus decreasing nonselective binding. After incubation, slides were rinsed 3 times with wash buffer reagents followed by Amplification steps I, II, and III. After Amplification III, slides were incubated in HRP, before they were incubated in TSA Plus Cy5, Cy3 or fluorescein at 1:1000 in TSA buffer (TSA^TM^ Plus System, Akoya Bioosciences). Following that slides were incubated in HRP blocker for 15 min and then rinsed 3 times with wash buffer reagent. Upon completion of the *in-situ* hybridization, slides immediately underwent immunohistochemistry for c-Fos (brain sections), NeuN and m-cherry (NDG sections).

### Immunohistochemistry

Immunohistochemistry was conducted immediately after *in situ* hybridization. Brain and NDG slides were rinsed 5 times in 50 mM potassium phosphate buffered saline (KPBS) for 5 min, and then blocked in 2% normal donkey serum and 0.2% Triton X-100 in KPBS to minimize nonspecific binding, before being incubated in the primary antibodies. For the NDG, slides were either incubated in Anti-NeuN or Anti-mCherry. For NeuN labelling, slides were incubated in the conjugated antibody A60 mouse anti-NeuN-AlexaFluor 488 (1:1000; MAB377X, EMD Millipore) for 2 h at room temperature. Slides were then rinsed in 50 mM KPBS allowed to air-dry, then cover-slipped. For mCherry labelling, to amplify the tdTomato signal, slides were incubated in primary antibody rabbit anti-mCherry (1:500; PA5-34974, Thermo Scientific) overnight at 4°C. Brain slides were incubated in the polyclonal rabbit anti-cFos (1:2000; RPCA-c-Fos, EnCor Biotechnology) overnight at 4°C. On the next day, slides were rinsed in rinsed in KPBS three times at room temperature. Following that, slides were incubated for 2 h in the secondary antibody donkey anti-rabbit Cy3 (711-165-152) purchased from Jackson ImmunoResearch and used at a 1:500 dilution. Slides were then rinsed with KPBS, incubated in DAPI (Advanced Cell Diagnostics, Newark, CA) for 30 s, rinsed again in KPBS, and allowed to air dried overnight. Slides were then cover slipped with ProLong^TM^ Gold antifade mounting medium before being imaged.

### Image capture

Images were captured and processed using a laser-scanning confocal microscope (Nikon Instruments Inc. Melville, NY, USA). Large coronal scans captured at 10 x magnification throughout the aortic arch and stomach (whole-mount), NDG (whole-mount LNG and RNG), and hindbrain (coronal sections) were acquired to assess the expression of tdTomato and GFP^3^. Regions with high expression of tdTomato and GFP were then imaged at 20 x magnification and z-stacks through sections were acquired.

For IHC and *in situ* hybridization in the brain and NDG, z-stacks of the proteins and mRNAs of interest were captured at 20 x or 40 x magnification. An average of 20 optical sections were collected per z-stack (0.5 µm between z-steps). For i*n situ* hybridization, sections hybridized with the probes of interest were used to determine the exposure time and image processing required to provide optimal visualization of RNA signal. As described in detail previously^8^, these same parameters were then used to assess background fluorescence in sections hybridized with the negative control probe (DapB). This allows for distinguishing bright positive control puncta (indicative of individual mRNA transcripts) from low-intensity background puncta. For evaluation of c-Fos-positive nuclei, the exposure time for the c-Fos channel was kept the same for all samples within a cohort. All representative photomicrographs were then prepared using FIJI^84^ and any adjustments to settings were kept consistent within a cohort.

### Quantification of RNAscope *in situ* hybridization and immunohistochemistry

Quantification of mRNA(s), proteins (c-Fos, NeuN and mCherry) and their colocalization was done on QuPath software. Maximum projection images from z-stacks of regions of interest (ROIs) of brain sections and NDG were quantified for mRNA expression and co-localizations with the different protein markers. For evaluation of percentage of c-Fos, NeuN, and m-Cherry cells containing mRNAs of interest, cells were considered to contain or colocalize to mRNA if at least three visible transcripts or puncta dots were observed with the volume of the protein markers. The mean number/percentages per section were determined by averaging the values across sections for each individual animal and then by calculating the mean across the group.

### Quantification of duodenal intraganglionic laminar endings (IGLEs)

Whole mount duodenum samples were prepared and analyzed as previously described^85^. Briefly, maximum intensity projections of confocal z-stack images were acquired from both tdTomato positive sensory afferent fibers and a separate background channel identifying muscle layers, ganglia, villi, and other tissue types. Quantitative analysis was performed using a 1 mm^2^ grid to determine IGLE density. IGLEs were identified as 1) a distinct cluster of terminal puncta, 2) the terminal portion of a single axon, and 3) overlapping ganglia identified on the background channel.

### Statistics

Data were analyzed and graphed using Prism 9 for Windows (GraphPad Software, Inc. La Jolla, CA). The number of intraganglionic laminar endings (IGLEs) per mm duodenum were analyzed using a 2-way analysis of variance (ANOVA) with Šídák’s multiple comparisons tests. Patterns of neuronal activation were analyzed using unpaired one-tailed t-tests. Acute and chronic CNO administration calorimetry, telemetry and food intake data were analyzed using either 2-way repeated measures ANOVAs or mixed effects models where appropriate. For acute administration, comparisons between Gq-NDG^Oxtr^ -CNO and Gq-saline, Con-CNO and Con-Saline data were made. For chronic CNO administration, CNO injection data were compared to Baseline and No Injection data within treatment group. When significant treatment or treatment x time interactions were observed, Dunnett’s multiple comparisons tests were performed. Anxiety-like behaviors (EPM, OF and LDB tests) and sociability were evaluated using unpaired two-tailed student t-tests. Conditioned taste avoidance was analyzed using 2-way RM ANOVA with Šídák’s multiple comparisons tests. Corticosterone responses to CNO administration were analyzed using 2-way RM ANOVA with Fisher’s LSD post-hoc tests. Differences in patterns of neuronal activation were analyzed using unpaired one-tailed t-tests. Detailed statistical results are provided as a supplementary table.

## Supporting information

Statistical Results Table 1

---

**Supplemental Figure 1.**
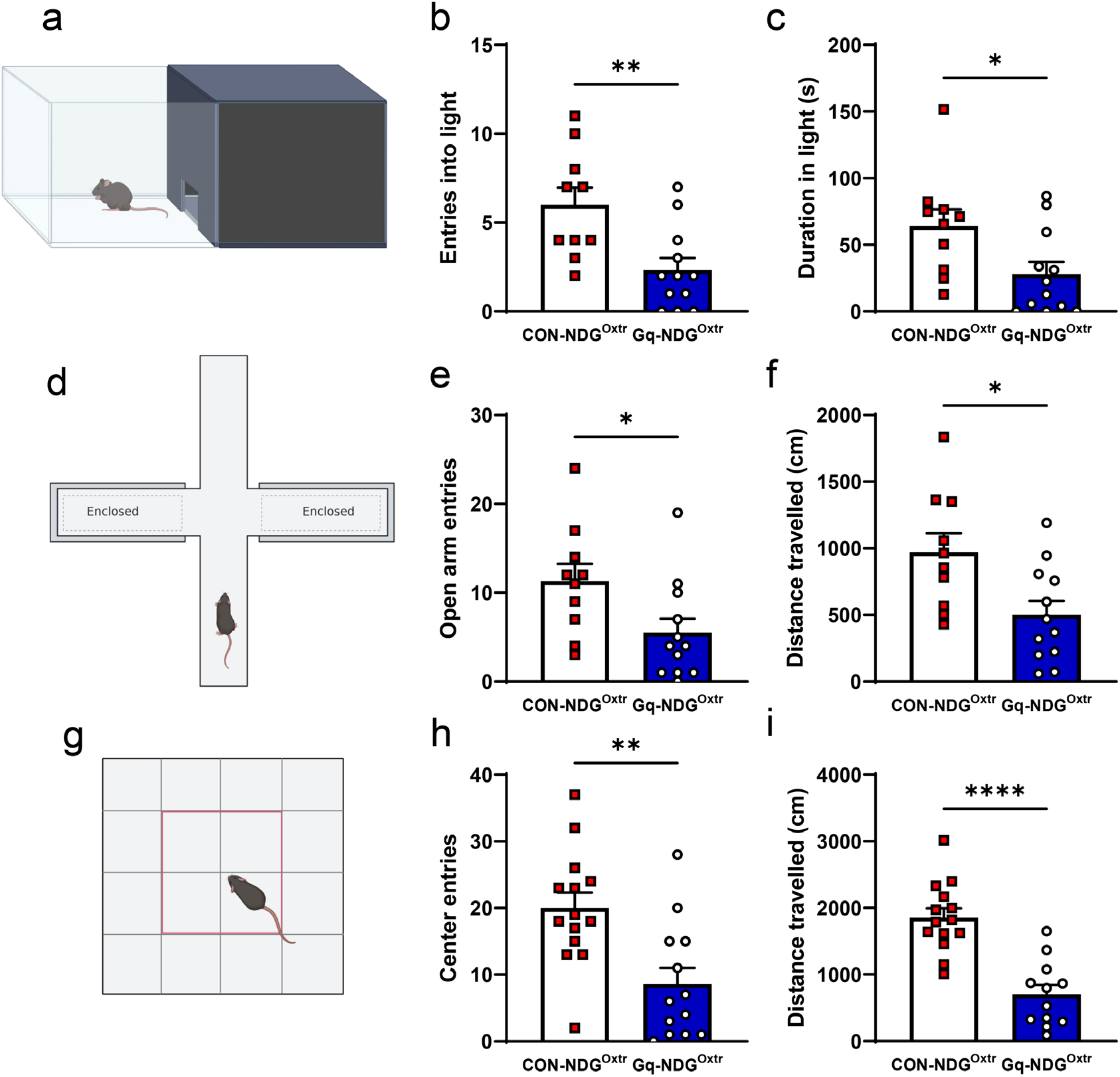
Acute activation of NDG^Oxtr^ induces anxiety-like behavior and reduces locomotor activity. **a,d,g** are schematics of the light dark box (LDB), elevated plus maze (EPM) and open field arena (OFA), respectively. Gq-NDG^Oxtr^ made fewer entries into the light **(b)** and spent less time in the light side (c) of the LDB. Gq-NDG^Oxtr^ mice made fewer entries **(e)** and travelled less in the EPM **(f)**. Gq-NDG^Oxtr^ made fewer center entries (g) and travelled less in the OFA (h). Data shown as mean ± s.e.m., n= 10-14 CON-NDG^Oxtr^, 12-13 Gq-NDG^Oxtr^. Unpaired one-tailed t-tests. *p<0.05, **p<0.01, ****p<0.0001. Schematics (a,d,g) made with Biorender.com.

**Supplemental Figure 2.**
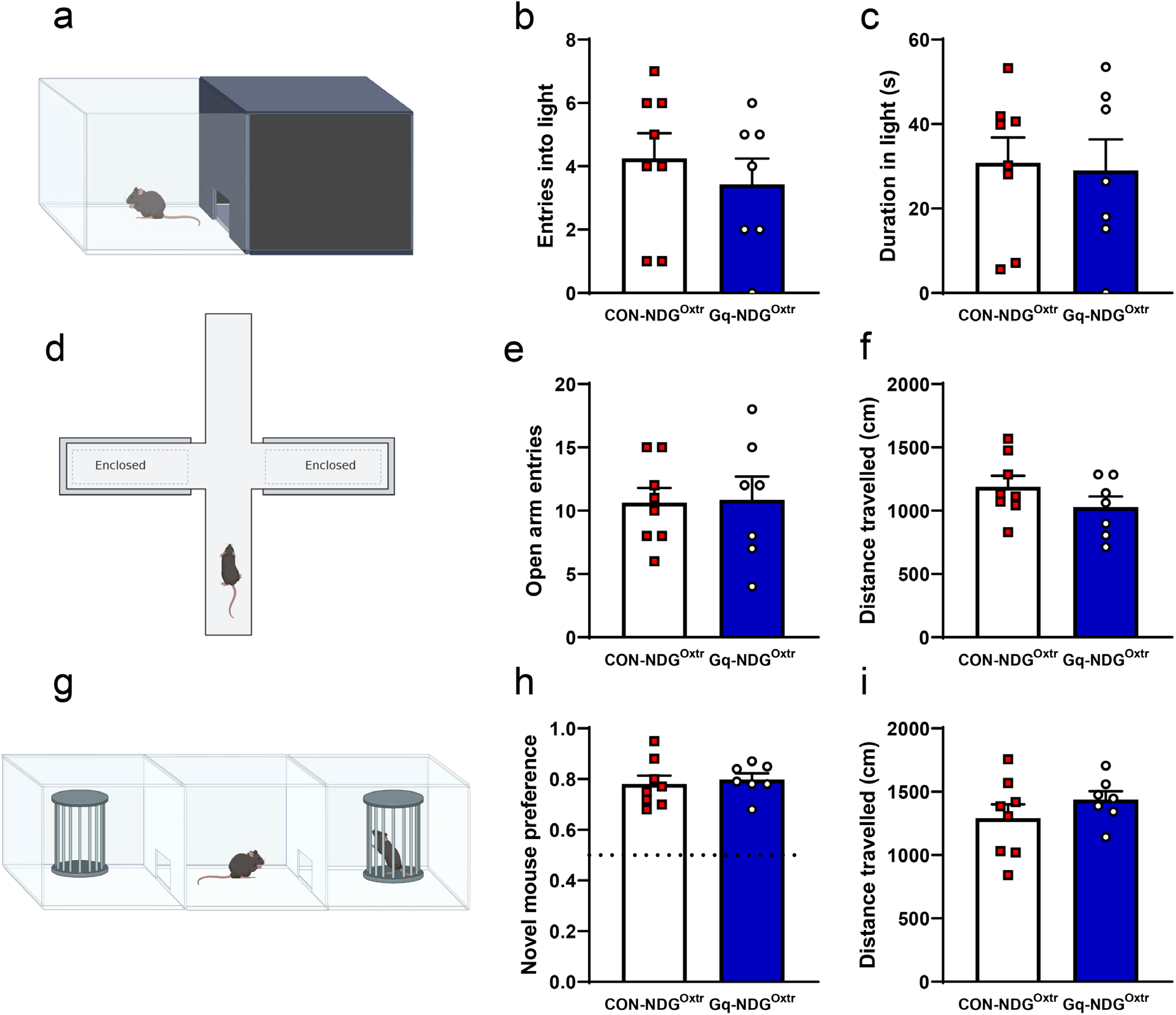
Chronic activation of NDG^Oxtr^ does not induce anxiety-like behavior or reduce social preference. a,d,g are schematics of the light dark box (LDB), elevated plus maze (EPM) and social interaction chamber, respectively. Entries into (b) and durations of time (c) spent in the light side of the LDB did not differ between groups. Open arm entries (e) and distance travelled (f)in the EPM did not differ between groups. Chronic activation of NDG^Oxtr^ did not affect preference investigating a novel mouse (h) or distanced travelled (i) in the 3-chamber social interaction test. Data shown as mean ± s.e.m., n= 8 CON-NDG^Oxtr^, 7 Gq-NDG^Oxtr^. Unpaired, one-tailed t-tests. Schematics (a,d,g) made with Biorender.com.

**Supplemental Figure 3.**
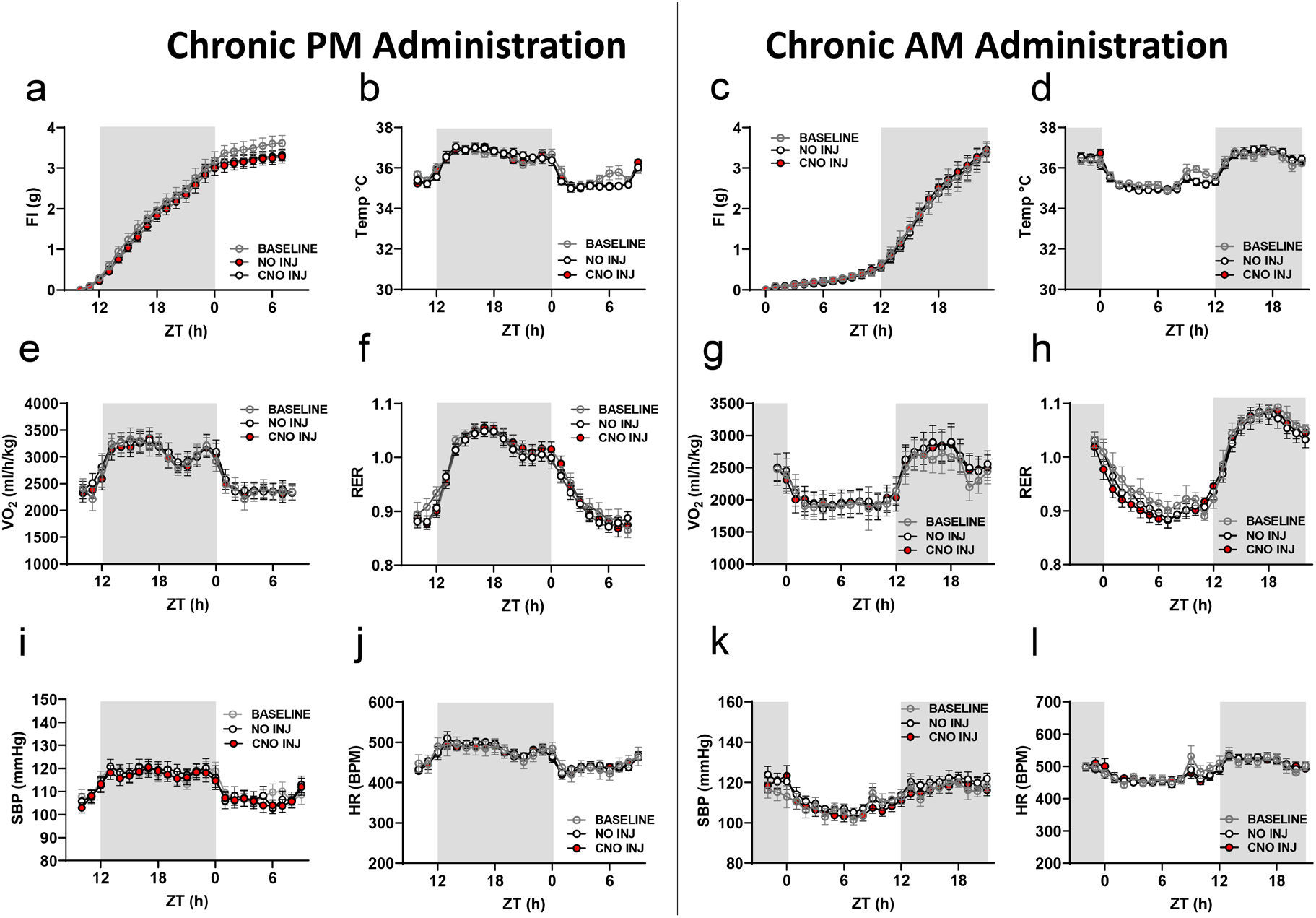
Chronic administration of CNO (0.3 mg/kg, i.p.) to CON-NDG^Oxtr^ (lacking Gq DREADDs) has no effect on food intake (a,c), core body temperature (b,d), oxygen consumption (e,g) or RER (f,h), SBP (i,k) or HR (j,l) when given 1h prior to onset of dark phase (left panel) or light phase (right panel). CON-NDG^Oxtr^ data are averaged between baseline (2 days), no injection (5 days) and CNO injection days (6 days). All figures express mean ± s.e.m. n= 15 CON (a,e,f), 7 CON (b,c,d,g,h,I,j,k,l)

## Acknowledgements

Funding sources NIH grants HL150750 (EGK), AT012142 (EGK, AdK, GdL).

## Author contributions

Karen A. Scott (study design, study implementation, data analysis, wrote the manuscript), Yalun Tan (study design, study implementation, data analysis), Dominique N. Johnson (study implementation, data analysis), Khalid Elsaafien (study implementation, data analysis, wrote the manuscript), Caitlin Baumer-Harrison (study implementation), Sophia A. Eikenberry (study implementation), Jessica M. Sa (study implementation), Guillaume de Lartigue (study design, study implementation, wrote the manuscript), Annette D. de Kloet (study design, study implementation, data analysis, wrote the manuscript), & Eric G. Krause (study design, study implementation, data analysis, wrote the manuscript).

## Data Availability Statement

All data supporting the findings of this study are available within the paper and its supplementary Information. Additional data are available from the corresponding author upon reasonable request.

