## Supplementary material for "Mechanosensation of the heart and gut elicits hypometabolism and vigilance in mice": Statistical Results Table 1

| Figure | Part | | Sample size | Test | P-value, F- value, df, t-values |
|---|---|---|---|---|---|
| 1 | k | IGLEs/mm duodenum | n= 3 males, 3 females | Two-way ANOVA, Šídák's multiple comparisons | Sex F (1,60)= 75.67, P<0.0001; Distance F (14, 60)= 12.37, P<0.0001; Sex x Distance F (14,60)= 8.694, P<0.0001 |

| Figure | Part | | Sample size | Test | P-value, F- value, df, t-values |
|---|---|---|---|---|---|
| 3 | b | Food Intake | n= 12 CON-NDG$^{Oxtr}$, n= 22 Gq-NDG$^{Oxtr}$ | Two-way RM ANOVA followed with Dunnett's multiple comparisons | Time F (19,1134)=364.4, P<0.0001; Treatment F (3,63)=18.20, P<0.0001; Time x Treatment F (54,1134)=7.408, P<0.0001 |
| 3 | c | VO2 | n= 12 CON-NDG$^{Oxtr}$, n= 22 Gq-NDG$^{Oxtr}$ | Mixed effects analysis with Dunnett's multiple comparisons | Time F (5.510, 331.2)= 25.02, P<0.0001; Treatment F (3,63)=12.72, P<0.0001; Time x Treatment F (54, 1082)= 7.305, P<0.0001 |
| 3 | d | RER | n= 12 CON-NDG$^{Oxtr}$, n= 22 Gq-NDG$^{Oxtr}$ | Mixed effects analysis with Dunnett's multiple comparisons | Time F(3.744, 225.7)= 27.31, P<0.0001; Treatment F (3, 64)= 12.79, P<0.0001; Time x Treatment F (54, 1085)= 11.72, P<0.0001 |
| 3 | e | Temp | n= 4 CON-NDG$^{Oxtr}$, n= 5 Gq-NDG$^{Oxtr}$ | Two-way RM ANOVA followed with Dunnett's multiple comparisons | Time F(18, 252)= 7.435, P<0.0001; Treatment F (3, 14)= 3.960, P= 0.0309; Time x Treatment F (54, 252)= 8.044, P<0.0001 |
| 3 | f | Systolic Blood Pressure | n= 4 CON-NDG$^{Oxtr}$, n= 5 Gq-NDG$^{Oxtr}$ | Two-way RM ANOVA followed with Dunnett's multiple comparisons | Time F(18, 252)= 12.09, P<0.0001; Treatment F (3, 14)= 0.3009, P= 0.8242; Time x Treatment F (54, 252)= 2.896, P<0.0001 |
| 3 | g | Heart Rate | n= 4 CON-NDG$^{Oxtr}$, n= 5 Gq-NDG$^{Oxtr}$ | Two-way RM ANOVA, Dunnett's multiple comparisons | Time F(18, 252)= 12.25, P<0.0001; Treatment F (3, 14)= 9.495, P= 0.0011; Time x Treatment F (54, 252)= 6.817, P<0.0001 |
| 3 | h | Water Intake | n= 12 CON-NDG$^{Oxtr}$, n= 22 Gq-NDG$^{Oxtr}$ | Two-way RM ANOVA, Dunnett's multiple comparisons | Time F (1.134, 71.42)=345.7, P<0.0001; Treatment F (3,63)=11.70, P<0.0001; Time x Treatment F (54,1134)=5.343, P<0.0001 |
| 3 | j | total fos AP | n= 4 CON-NDG$^{Oxtr}$, n= 4 Gq-NDG$^{Oxtr}$ | Unpaired one-way t-test | p= 0.0183, t= 2.677, df= 6 |
| 3 | j | total fos NTS | n= 4 CON-NDG$^{Oxtr}$, n= 4 Gq-NDG$^{Oxtr}$ | Unpaired one-way t-test | p= 0.0018, t= 4.618, df= 6 |
| 3 | j | GIPRr+ Fos+/GIPR+ AP | n= 4 CON-NDG$^{Oxtr}$, n= 4 Gq-NDG$^{Oxtr}$ | Unpaired one-way t-test | p= 0.0684, t= 1.716, df= 6 |

| Figure | Part | | Sample size | Test | P-value, F- value, df, t-values |
|---|---|---|---|---|---|
| 3 | j | GIPR+ Fos+/GIPR+ NTS | n= 4 CON-NDG$^{Oxtr}$, n= 4 Gq-NDG$^{Oxtr}$ | Unpaired one-way t-test | p= 0.2977, t= 0.5604, df= 6 |
| 3 | j | VGAT+Fos+/VGAT+ AP | n= 4 CON-NDG$^{Oxtr}$, n= 4 Gq-NDG$^{Oxtr}$ | Unpaired one-way t-test | p= 0.0257, t= 2.427, df= 6 |
| 3 | j | VGAT+Fos+/VGAT+ NTS | n= 4 CON-NDG$^{Oxtr}$, n= 4 Gq-NDG$^{Oxtr}$ | Unpaired one-way t-test | p= 0.0007, t= 5.607, df= 6 |
| 3 | j | GIPR+Fos+VGAT+/GIPR +VGAT+ AP | n= 4 CON-NDG$^{Oxtr}$, n= 4 Gq-NDG$^{Oxtr}$ | Unpaired one-way t-test | p= 0.0053, t= 3.651, df= 6 |
| 3 | j | GIPR+Fos+VGAT+/GIPR +VGAT+ NTS | n= 4 CON-NDG$^{Oxtr}$, n= 4 Gq-NDG$^{Oxtr}$ | Unpaired one-way t-test | p=0.0087, t= 3.252, df= 6 |
| 3 | l | total c-Fos PBN | n= 5 CON-NDG$^{Oxtr}$, n= 5 Gq-NDG$^{Oxtr}$ | Unpaired one-way t-test | p= 0.0479, t= 1.887, df= 8 |
| 3 | l | CALCA+Fos+/CALCA+ PBN | n= 5 CON-NDG$^{Oxtr}$, n= 4 Gq-NDG$^{Oxtr}$ | Unpaired one-way t-test | p<0.0001, t= 7.692, df= 7 |
| 3 | m | Flavor Preference | n= 6 CON-NDG$^{Oxtr}$, n= 7 Gq-NDG$^{Oxtr}$ | Two-way RM ANOVA, Šídák's multiple comparisons | Injection F(1, 11)= 5.189, P<0.0437; Treatment F (1, 11)= 5.422, P= 0.0400; Time x Treatment F (1, 11)= 17.53, P<0.0015 |
| 3 | o | Total c-Fos PVN | n= 4 CON-NDG$^{Oxtr}$, n= 4 Gq-NDG$^{Oxtr}$ | Unpaired one-way t-test | p= 0.0044, t= 3.807, df= 6 |
| 3 | o | CRH+ c-Fos+/CRH+pvn | n= 4 CON-NDG$^{Oxtr}$, n= 4 Gq-NDG$^{Oxtr}$ | Unpaired one-way t-test | p= 0.0031, t= 4.111, df= 6 |
| 3 | p | Plasma Corticosterone | n= 5 CON-NDG$^{Oxtr}$, n= 5 Gq-NDG$^{Oxtr}$ | Two-way RM ANOVA, Fisher's LSD | Time F(2.184, 17.47)= 16.53, P<0.0001; Treatment F (1, 8)= 18.82, P= 0.0025; Time x Treatment F (4, 32)= 7.927, P= 0.0001 |

| Figure | Part | | Sample size | Test | P-value, F- value, df, t-values |
|---|---|---|---|---|---|
| 4 | a | PM Food Intake | n= 13 Gq-NDG$^{Oxtr}$ | Two-way RM ANOVA, Dunnett's multiple comparisons | Time F (21,756)= 5.880, P<0.0001; Treatment F (2, 36)= 10.61, P<0.0001; Time x Treatment F (42,756)= 5.880, P<0.0001 |
| 4 | b | PM Temp | n= 8 Gq-NDG$^{Oxtr}$ | Two-way RM ANOVA, Dunnett's multiple comparisons | Time F (23, 483)= 11.08 , P<0.0001; Treatment F (2, 21)= 14.32, P=0.0001; Time x Treatment F (46, 483)= 20.61 , P<0.0001 |
| 4 | c | AM Food Intake | n= 8 Gq-NDG$^{Oxtr}$ | Mixed-effects analysis, Dunnett's multiple comparisons | Time F (1.372, 28.70)= 280.8, P<0.0001; Treatment F (2,21)= 6.808, P<0.0053; Time x Treatment F (46, 481)= 2.388 , P<0.0001 |
| 4 | d | AM Temp | n= 8 Gq-NDG$^{Oxtr}$ | Two-way RM ANOVA, Dunnett's multiple comparisons | Time F (2.752, 57.79)= 53.18, P<0.0001; Treatment F (2, 21)= 7.793, P=0.0029; Time x Treatment F (46, 483)= 8.032, P<0.0001 |
| 4 | e | PM VO2 | n= 13 Gq-NDG$^{Oxtr}$ | Mixed-effects analysis followed with Dunnett's multiple comparisons | Time F (5.617, 197.1)= 37.57 , P<0.0001; Treatment F (2, 36)= , P= 0.3764; Time x Treatment F (44, 772)= 11.33, P<0.0001 |
| 4 | f | PM RER | n= 13 Gq-NDG$^{Oxtr}$ | Mixed-effects analysis, Dunnett's multiple comparisons | Time F (21, 732)= 31.52, P<0.0001; Treatment F (2, 36)= 4.393, P=0.196; Time x Treatment F (42,732)= 18.38, P<0.0001 |
| 4 | g | AM VO2 | n= 8 Gq-NDG$^{Oxtr}$ | Mixed-effects analysis, Dunnett's multiple comparisons | Time F (5.776, 120.5)= 49.18 , P<0.0001; Treatment F (2, 21)= 0.4999, P=0.6136; Time x Treatment F (46, 480)= 2.858, P<0.0001 |
| 4 | h | AM RER | n= 8 Gq-NDG$^{Oxtr}$ | Mixed-effects analysis, Dunnett's multiple comparisons | Time F (3.616, 75.47)= 75.74, P<0.0001; Treatment F (2, 21)= 4.345 , P=0.0264; Time x Treatment F (46, 480)= 3.926, P<0.0001 |

| Figure | Part | | Sample size | Test | P-value, F- value, df, t-values |
|---|---|---|---|---|---|
| 4 | i | PM SBP | n= 8 Gq-NDG$^{Oxtr}$ | Two-way RM ANOVA, Dunnett's multiple comparisons | Time F (6.018, 126.4)= 10.23 , P<0.0001; Treatment F (2, 21)= 0.5895 , P=0.5635; Time x Treatment F (46,483)= 4.701, P<0.0001 |
| 4 | j | PM HR | n= 8 Gq-NDG$^{Oxtr}$ | Two-way RM ANOVA, Dunnett's multiple comparisons | Time F (4.374, 91.85)= 7.973 , P<0.0001; Treatment F (2, 21)= 11.74 , P= 0.0004; Time x Treatment F (46, 483)= 19.88 , P<0.0001 |
| 4 | k | AM SBP | n= 8 Gq-NDG$^{Oxtr}$ | Two-way RM ANOVA, Dunnett's multiple comparisons | Time F (3.617, 75.95)= 20.34 , P<0.0001; Treatment F (2,21)= 0.3491, P= 0.7094; Time x Treatment F (46, 483)= 1.739, P= 0.0026 |
| 4 | l | AM HR | n= 8 Gq-NDG$^{Oxtr}$ | Two-way RM ANOVA, Dunnett's multiple comparisons | Time F (6.122, 128.6)= 50.45, P<0.0001; Treatment F (2, 21)= 12.25, P= 0.0003; Time x Treatment F (46, 483)= 8.336, P<0.0001 |
| 4 | m | PM Body Weight | n= 17 CON-NDG$^{Oxtr}$, r | Mixed-effects analysis, Šídák's multiple comparisons | Time F (11, 308)= 12.34, P<0.0001; Treatment F (1, 28)= 34.60, P<0.0001; Time x Treatment F (11, 308)= 6.393, P<0.0001 |
| 4 | n | AM Body Weight | n= 11 CON-NDG$^{Oxtr}$, r | Mixed-effects analysis, Šídák's multiple comparisons | Time F (6.401, 108.3)= 8.097, P<0.0001; Treatment F (1, 17)= 22.16, P= 0.0002; Time x Treatment F (12, 203)= 2.887, P= 0.0011 |

| Figure | Part | | Sample size | Test | P-value, F- value, df, t-values |
| --- | --- | --- | --- | --- | --- |
| S1 | b | LDB Light entries | n= 10 CON-$NDG^{Oxtr}$, n= 12 Gq-$NDG^{Oxtr}$ | Unpaired two-tailed t-test | p= 0.0044, t= 3.206, df= 20 |
| S1 | c | LDB Light duration | n= 10 CON-$NDG^{Oxtr}$, n= 12 Gq-$NDG^{Oxtr}$ | Unpaired two-tailed t-test | p= 0.0129, t= 2.408, df= 20 |
| S1 | e | EPM Open arm entries | n= 10 CON-$NDG^{Oxtr}$, n= 12 Gq-$NDG^{Oxtr}$ | Unpaired two-tailed t-test | p= 0.0157, t= 2.314, df= 20 |
| S1 | f | EPM distance | n= 10 CON-$NDG^{Oxtr}$, n= 12 Gq-$NDG^{Oxtr}$ | Unpaired two-tailed t-test | p= 0.0127, t= 2.737, df= 20 |
| S1 | h | OFA center entries | n= 14 CON-$NDG^{Oxtr}$, n= 13 Gq-$NDG^{Oxtr}$ | Unpaired two-tailed t-test | p= 0.0022, t= 3.417, df= 25 |
| S1 | i | OFA distance | n= 14 CON-$NDG^{Oxtr}$, n= 13 Gq-$NDG^{Oxtr}$ | Unpaired two-tailed t-test | p<0.0001, t= 5.788, df= 24 |

| Figure | Part | | Sample size | Test | P-value, F- value, df, t-values |
|---|---|---|---|---|---|
| S2 | b | LDB Light entries | n= 8 CON-NDG$^{Oxtr}$, n= 7 Gq-NDG$^{Oxtr}$ | Unpaired two-tailed t-test | p= 0.2422, t= 0.7198, df= 13 |
| S2 | c | LDB Light duration | n= 8 CON-NDG$^{Oxtr}$, n= 7 Gq-NDG$^{Oxtr}$ | Unpaired two-tailed t-test | p= 0.8509, t= 0.1918, df= 13 |
| S2 | e | EPM Open arm entries | n= 8 CON-NDG$^{Oxtr}$, n= 7 Gq-NDG$^{Oxtr}$ | Unpaired two-tailed t-test | p= 0.9142, t= 0.1099, df= 13 |
| S2 | f | EPM distance | n= 8 CON-NDG$^{Oxtr}$, n= 7 Gq-NDG$^{Oxtr}$ | Unpaired two-tailed t-test | p= 0.2043, t= 1.337, df= 13 |
| S2 | h | Mouse preference | n= 8 CON-NDG$^{Oxtr}$, n= 7 Gq-NDG$^{Oxtr}$ | Unpaired two-tailed t-test | p= 0.6850, t= 0.4149, df= 13 |
| S2 | i | Distance travelled | n= 8 CON-NDG$^{Oxtr}$, n= 7 Gq-NDG$^{Oxtr}$ | Unpaired two-tailed t-test | p= 0.2905, t= 1.102, df= 13 |

| Figure | Part | | Sample size | Test | P-value, F- value, df, t-values |
|---|---|---|---|---|---|
| S3 | a | PM Food Intake | n= 15 CON-$NDG^{Oxtr}$ | Mixed-effects analysis, Dunnett's multiple comparisons | Time F (2.430, 102.1)= 625.2 , P<0.0001; Treatment F (2,42)= 0.6727, P= 0.5158; Time x Treatment F (42, 882)= 0.5109, P= 0.9960 |
| S3 | b | PM Temp | n= 7 CON-$NDG^{Oxtr}$ | Mixed-effects analysis, Dunnett's multiple comparisons | Time F (4.551, 77.36)= 81.80 , P<0.0001; Treatment F (2,17)= 0.0.2157, P= 0.8081; Time x Treatment F (46, 391)= 1.633, P= 0.0076 |
| S3 | c | AM Food intake | n= 7 CON-$NDG^{Oxtr}$ | Two-way RM ANOVA, Dunnett's multiple comparisons | Time F (1.347, 24.24)= 295.4, P<0.0001; Treatment F (2, 18)= 0.07495, P= 0.9281; Time x Treatment F (46,414)= 0.07942, P>0.9999 |
| S3 | d | AM Temp | n= 7 CON-$NDG^{Oxtr}$ | Two-way RM ANOVA, Dunnett's multiple comparisons | Time F (4.984, 89.70)= 69.90, P<0.0001; Treatment F (2, 18)= 0.1290, P=0.8798; Time x Treatment F (46, 414)= 1.072, P= 0.6523 |
| S3 | e | PM $VO_2$ | n= 15 CON-$NDG^{Oxtr}$ | Mixed-effects analysis, Dunnett's multiple comparisons | Time F (6.349, 260.9)= 90.69, P<0.0001; Treatment F (2, 42)= 0.02596, P= 0.9744; Time x Treatment F (44, 904)= 0.6956, P= 0.9341 |
| S3 | f | PM RER | n= 15 CON-$NDG^{Oxtr}$ | Mixed-effects analysis, Dunnett's multiple comparisons | Time F (4.271, 175.3)= 139.6, P<0.0001; Treatment F (2, 42)= 0.1282, P=0.8800; Time x Treatment F (44, 903)= 0.5772, P= 0.9879 |
| S3 | g | AM $VO_2$ | n= 7 CON-$NDG^{Oxtr}$ | Mixed-effects analysis, Dunnett's multiple comparisons | Time F (5.637, 101.0)= 66.56, P<0.0001; Treatment F (2, 18)= 0.03491, P= 0.9658; Time x Treatment F (46, 412)= 0.6729; P= 0.9503 |
| S3 | h | AM RER | n= 7 CON-$NDG^{Oxtr}$ | Mixed-effects analysis, Dunnett's multiple comparisons | Time F (3.755, 67.27)= 108.2, P<0.0001; Treatment F (2, 18)= 0.4988, P= 0.6154; Time x Treatment F (46, 412)= 0.6736, P= 0.9499 |
| S3 | i | PM SBP | n= 6 CON-$NDG^{Oxtr}$ | Two-way RM ANOVA, Dunnett's multiple comparisons | Time F (5.997, 89.90)= 38.09, P<0.0001; Treatment F (2, 15)= 0.05188, P= 0.9496; Time x Treatment F (46, 345)= 0.9880, P= 0.4990 |

| Figure | Part | | Sample size | Test | P-value, F- value, df, t-values |
|---|---|---|---|---|---|
| S3 | j | PM HR | n= 7 CON-NDG$^{Oxtr}$ | Two-way RM ANOVA, Dunnett's multiple comparisons | Time F (4.788, 86.18)= 29.45, P<0.0001; Treatment F (2, 18)= 0.02076, P= 0.9795; Time x Treatment F (46, 414)= 0.7147, P= 0.9193 |
| S3 | k | AM SBP | n= 6 CON-NDG$^{Oxtr}$ | Two-way RM ANOVA, Dunnett's multiple comparisons | Time F (4.109, 61.63)= 30.60, P<0.0001; Treatment F (2, 15)= 0.3507, P= 0.7098; Time x Treatment (46, 345)= 1.233, P= 0.1527 |
| S3 | l | AM HR | n= 7 CON-NDG$^{Oxtr}$ | Two-way RM ANOVA, Dunnett's multiple comparisons | Time F (6.154, 110.8)= 28.58, P<0.0001; Treatment F (2, 18)= 0.03459, P= 0.9661; Time x Treatment F (46, 414)= 1.073, P=0.3512 |
